# Reward Bases: Instantaneous reward revaluation with temporal difference learning

**DOI:** 10.1101/2022.04.14.488361

**Authors:** Beren Millidge, Mark Walton, Rafal Bogacz

## Abstract

An influential theory posits that dopaminergic neurons in the mid-brain implement a model-free reinforcement learning algorithm based on temporal difference (TD) learning. A fundamental assumption of this model is that the reward function being optimized is fixed. However, for biological creatures the ‘reward function’ can fluctuate substantially over time depending on the internal physiological state of the animal. For instance, food is rewarding when you are hungry, but not when you are satiated. While a variety of experiments have demonstrated that animals can instantly adapt their behaviour when their internal physiological state changes, under current thinking this requires model-based planning since the standard model of TD learning requires retraining from scratch if the reward function changes. Here, we propose a novel and simple extension to TD learning that allows for the zero-shot (instantaneous) generalization to changing reward functions. Mathematically, we show that if we assume the reward function is a linear combination of *reward basis vectors*, and if we learn a value function for each reward basis using TD learning, then we can recover the true value function by a linear combination of these value function bases. This representational scheme allows instant and perfect generalization to any reward function in the span of the reward basis vectors as well as possesses a straightforward implementation in neural circuitry by parallelizing the standard circuitry required for TD learning. We demonstrate that our algorithm can also reproduce behavioural data on reward revaluation tasks, predict dopamine responses in the nucleus accumbens, as well as learn equally fast as successor representations while requiring much less memory.

## Introduction

An influential viewpoint in the cognitive neurosciences is that much reward-driven animal and human behaviour can be well explained through the lens of reinforcement learning (Dayan, 1993; Dayan & Daw, 2008; Schultz, 1998, 2002; Sutton & Barto, 2018) – a theory which emerged out of concepts of utility maximization in game theory and economics (Von Neumann & Morgenstern, 2007; Zeki, Goodenough, & Zak, 2004) combined with Pavlovian and Behaviourist notions of stimulus-response associations and conditioning (Averbeck & Costa, 2017; Medina, Repa, Mauk, & LeDoux, 2002; Mollick et al., 2020; Pavlov & Gantt, 1928; Skinner, 1985). Reinforcement learning theory assumes that agentic behaviour can be modelled as choosing actions in the world so as to maximize a reward function. Given a reward function, the field of reinforcement learning provides a large number of effective methods for learning or computing actions that reliably lead to a high reward (Bertsekas, 2012; Sutton & Barto, 2018).

Due to the computational success and mathematical elegance of the reinforcement learning paradigm, it has long been suspected that the brain may implement reinforcement learning algorithms internally. A key breakthrough came when it was demonstrated that dopaminergic neurons in the ventral tegmental area (VTA), which strongly project to the striatum in the basal ganglia, appear to compute reward prediction errors (Friston, Tononi, Reeke Jr, Sporns, & Edelman, 1994; Houk & Adams, 1995; Montague, Dayan, & Sejnowski, 1996; Schultz, Dayan, & Montague, 1997). Reward prediction errors (RPEs) are the difference between the reward predicted by the brain and the reward actually received. These RPEs are a crucial ingredient in temporal difference (TD) model-free reinforcement learning methods, and thus this finding was developed into the ‘standard model’ of the basal ganglia as directly implementing TD learning (Dayan & Balleine, 2002; Schultz, 2002).

TD learning aims to infer the *value function*, which is the discounted sum of future rewards, for each state an animal might encounter. While a seemingly complicated mathematical object, TD learning provides a simple and mathematically elegant ways to approximate this value function directly from experience using RPEs. Specifically, at each timestep, we can observe the current reward as well as make a prediction of the expected value of the state. From this, we can compute the RPE as the difference between the two and use it to adjust our estimated value function. As long as all states can be visited, over time this procedure will lead an unbiased estimate of the true value function (Bertsekas, 2019; Sutton & Barto, 2018).

However, an implicit assumption of model-free reinforcement learning algorithms, including TD learning, is that the reward function is fixed. This is because the value function is a cached estimate of future reward, and so if these future rewards change, then the value function becomes out of date. However, for understanding the behaviour of many biological organisms we *cannot* assume that the reward function is fixed since it appears to change adaptively based on internal states and beliefs. For instance, food is typically rewarding until you are full at which point it becomes neutral, or even aversive. The same is true of most other ‘intrinsically rewarding’ stimuli such as water, sugar, chocolate, and so on. These effects are general examples of what is known as the ‘hedonic treadmill’ in which rewarding stimuli become less rewarding the more they are obtained until they end up neutral or even aversive in character (Barto, 2013; Castro, Cole, & Berridge, 2015; Kelley, Baldo, & Pratt, 2005). Indeed, biological brains appear to have developed a complex set of machinery for manipulating reward functions dynamically. This differs substantially from classical reinforcement agents where the reward function is assumed to be completely fixed. For RL agents, the billionth sip of sugar juice is just as good as the first ^1^

Beyond purely phenomenological evidence for changing reward functions, there is also substantial evidence that animals *can*, in fact, flexibly adapt their behaviour in the face of changes to the reward function. Moreover, animals often exhibit *zero-shot generalization*, meaning that if the reward function changes, they can instantaneously adapt their behaviour without having to ever experience the new reward function or have any new interactions with the environment at all. A compelling demonstrations of this capability come from a series of experiments (Robinson & Berridge, 2013) involving rats which learn to associate two cues with receiving either pleasant sugar juice or extremely unpleasant salt water with the salt concentration level of the Dead Sea. The rats all very quickly learn to approach the lever associated with the juice and avoid the lever associated with the salt solution. Then, once the rats have learned this task, they are placed in a physiological state which mimics the brain chemistry of salt deprivation. Importantly, this is a completely novel sensation for them, since during their entire life (as laboratory rats) they have never previously been salt deprived. When, in this virtual salt-deprived state, the rats are then given access to the same pair of levers, they immediately approach the lever which is associated with the salt water and begin making orofacial ‘liking’ behaviours. This occurs even if the salt-associated lever is presented *in extinction*, meaning that the animal never experiences the salt in the salt deprived state. Essentially, the rats appear able to instantaneously switch their reactions and responses to the sea salt lever, based purely on their internal physiological state, without ever having experienced the salty water being rewarding, nor even having experienced the salt-deprived state before.

This cannot be explained through standard reinforcement learning algorithms such as TD learning which require an experience of reward to update the value function estimate. TD learning would predict that the salt-deprived animals would still avoid the salty lever until the salt solution is delivered which would give them an unexpectedly positive reward, and would slowly induce them to approach that lever more and more often until it becomes the primary lever to be approached.^2^. Beyond this classic demonstration, there is a wide range of literature arguing that Pavlovian learned associations appear to be dynamically responsive not just to experienced rewards but to internal physiological states (DiFeliceantonio, Mabrouk, Kennedy, & Berridge, 2012; Krashes et al., 2009; Peciña & Berridge, 2013; Smith, Berridge, & Aldridge, 2011), as well as sensitive to the precise identities of the stimuli that have been learned (Dickinson, 1986; Holland, Lasseter, & Agarwal, 2008; Rescorla, 1988).

There have been several explanations proposed for this zero-shot reward revaluation capability in the literature. Dayan and Berridge (2014) proposed that the ability to achieve zero-shot reward revaluation is due to model-based planning. If the planner has access to the true internal reward model, and can use it to query the reward of arbitrary imagined states during planning, then zero-shot reward revaluation is, of course, straightforward. There is a large literature demonstrating that humans and other animals often employ hybrid strategies comprising both model-free and model-based strategies (Dolan & Dayan, 2013; Doll, Simon, & Daw, 2012; O’Doherty, Lee, & McNamee, 2015), and often flexibly switch between them based upon the importance of the task and the computational load (Daw & Dayan, 2014; Gläscher, Daw, Dayan, & O’Doherty, 2010), since model-based planning is generally considered more ‘computationally intensive’ for the brain, and thus relied on only when the model-free system cannot handle the requisite decision-making. Another approach is to use successor representations (Dayan, 1993) which represent a ‘successor matrix’ of discounted state occupancies allowing value functions to be estimated for any given reward function. A final approach, in the context of homeostatically driven changes in reward function, is to directly augment the reward term with a ‘homeostatic’ term which modulates the predicted reward according to homeostatic factors (Zhang, Berridge, Tindell, Smith, & Aldridge, 2009). This model can solve problems which only require a single step, such as the sea-salt experiments, where accurately estimating the reward is sufficient to solve the task, but does not suffice for flexibly changing behaviour when the goal requires multiple steps to achieve – since the long-term value function is not modulated.

There is strong evidence that the basal ganglia system is a core component in reward-based action selection (Daw & Dayan, 2014; Dayan & Balleine, 2002; Dayan & Berridge, 2014; Schultz, 2016; Schultz et al., 1997; Sesack & Grace, 2010; S. C. Tanaka et al., 2016), and is linked to brain systems controlling physiological state such as the hypothalamus (Godfrey & Borgland, 2019; Kelley, Baldo, Pratt, & Will, 2005; Morales & Berridge, 2020; O’Connor et al., 2015) such that there is a tight closed loop between hypothalamus and VTA dopamine neurons. Moreover, there is evidence that the basal ganglia model-free system may be relied on *more* than the model-based system when physiological needs are pressing (van Swieten, Bogacz, & Manohar, 2021), as well as that dorsomedial striatal lesions abolish flexible reward devaluation behaviour. However, the standard computational model for the function of the basal ganglia – TD learning – cannot handle changing reward functions without retraining from scratch – a property which is completely infeasible for the brain where the ‘reward function’ often varies continuously. Moreover, there is very little evidence that the basal ganglia possesses the requisite microcircuit architecture to directly implement model-based or successor representation based methods ^3^. There is thus a fundamental difficulty with the standard theory: in practice, the basal ganglia *must* be able to flexible adapt to changing reward functions if it is to function at all, yet TD learning, the algorithm it supposedly implements, simply cannot do this.

In this paper, we propose a potential solution to this difficulty. We derive a novel algorithm which we call *reward bases* that allows for zero-shot generalization to a changing reward function while requiring only a slight extension of the TD learning scheme. Specifically, our approach assumes that the reward function can be expressed as a linear combination of what we call *reward basis vectors*. We show that if we learn a separate value function for each reward basis vector, using normal TD learning, then we can combine these value function bases together flexibly to dynamically compute the true value function given a changing reward function. Our approach offers comparable zero-shot reward revaluation capabilities to successor representations but requires much less memory. Although it is not quite as expressive (meaning it cannot represent all possible reward functions) as model-based methods, it is computationally cheaper and provides a very accurate approximation in practice. We also present a more biologically plausible version of our model which directly modulates the prediction error signals of the dopamine neurons. This modified model allows us to successfully fit neural data of dopamine release in paradigms which cannot be explained by TD learning models and also provides a simple and natural implementation in the circuitry of the basal ganglia as a straightforward extension of the standard model of (Schultz, 1998). This paper thus makes two core contributions – first a novel computational algorithm in reinforcement learning which achieves the same generalization capabilities as successor representations with a smaller computational cost. Secondly, a biologically plausible implementation which could be implemented in the circuitry of the basal ganglia and which can match neural data.

The rest of the paper will be structured as follows. We first review the TD learning algorithm. Next, we present the mathematical details of our theory of reward bases. Then we propose a potential neural architecture for how this approach could be implemented in a model-free reinforcement learning circuit by modifying the original ideas of Montague et al. (1996); Schultz et al. (1997), and discuss further extensions of our theory in terms of directly modulating the RPEs by physiological state. Subsequently, we investigate how our approach performs in several simulated decision tasks with reward revaluation, including a simulated version of the Dead Sea salt paradigm (Robinson & Berridge, 2013) and demonstrate that our method can perform instant zero-shot reward revaluation unlike TD learning. We demonstrate that our model can reproduce experimentally observed dopaminergic response patterns in the nucleus accumbens. Crucially, we show that reward basis can adapt choices according to the internal state also for multi-step policies, which is not the case in other models of reward revaluation in the literature, as well as learns equally fast as the successor representations on a more complex simulated task. Finally, we discuss the evidence for and against the proposed model, as well as make novel experimental predictions.

### Temporal difference learning

We formalize our agent’s environment as a Markov Decision Process (*𝒳, 𝒜, ℛ, T, γ*) comprising states **x** ∈ *𝒳*, actions **a** ∈ *𝒜*, rewards *r* ∈ *ℛ*, a finite time-horizon *T*, and a scalar discount rate 0 ≤ *γ* ≤ 1. We denote vectors in **bold**. For simplicity, we consider discrete states, but the generalization to continuous state spaces with function approximation methods is relatively straightforward. The state of the environment **x** is represented as a one-hot vector of values for each discrete state. The agent can emit discrete actions **a** (where actions are represented as a one-hot vector over the action space) which change the transition dynamics of the environment. We consider the environmental dynamics to be stochastic with transition dynamics *p*(**x**_*t*_|**x**_*t*−1_, **a**_*t*−1_) which denotes the probability of moving from one state to another given a certain action **a**_*t*−1_. A *policy π* is defined as a function that maps a state vector **x** to an action vector **a**. The agent additionally computes a reward function *r*(**x**) which outputs a deterministic scalar reward for a given state.

The agents’ goal is to maximize the sum of discounted future rewards up until a time horizon *T* ^4^. This discounted sum is often called the return: 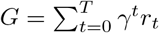, In order to maximize this return, a key quantity is the *value function* of a state, defined as the (scalar) expected return given the current policy at a specific timestep, starting from that state. The value function satisfies the recursive Bellman equation,

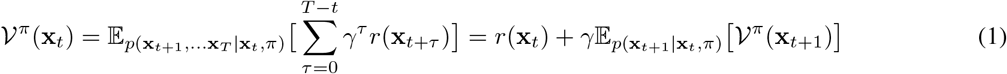

Crucially, this recursive relationship can be exploited to let us *estimate* the value function using only single transitions. This can be done by estimating the value function *𝒱*^*π*^ (**x**_*t*_) for each state and updating these estimates according to the difference between the estimated value function before receiving the reward and a newly estimated value function based on Equation 1. Due to the discrete state-space, we can define the transition operator *𝒯 ^π^* such that 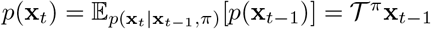. In practice, *𝒯 ^π^* is a | *𝒳* | × | *𝒳* | matrix (where | · | denotes the size of a set) which denotes the probability of transitioning from one state to another given a specific policy (or action), and the transition operator can be implemented as a matrix-multiplication in discrete state spaces. Using this new notation, we can rewrite Equation 1 for the value function in a simpler way,

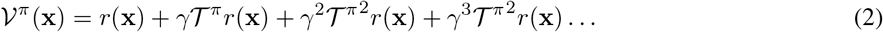

While the Bellman recursion can be expressed as,

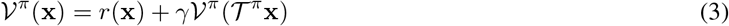

where **x***′* is the next state that occurs given the previous state and action. We can compute a RPE *δ*(**x**) as the difference between the estimated value function and that computed by the Bellman recursion, and then update our estimated value function in proportion to the RPE

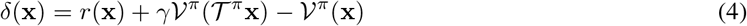

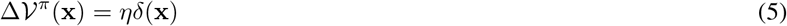

where *η* is a scalar learning rate. This algorithm is known as TD learning and converges to the true value function if all states are visited a sufficient number of times (Sutton & Barto, 2018).

## Results

The fundamental idea presented in this paper is that instead of considering a single scalar reward function which the animal computes and maximizes, we instead suppose that the reward function is composed of a *linear combination* of component reward functions which, in analogy to coordinate bases in linear algebra, we call *reward bases*. Given these reward bases, we can then weight them in different ways to construct a variety of different total reward functions. The key insight is that if the total reward function is comprised of a linear combination of reward bases, then the total value function can also be computed as a linear combination of the *value functions* of each individual reward basis. This enables classical TD learning to be performed for each reward basis independently to compute a value basis for each reward basis. Then, given a specific weighting of the reward bases, the total value function can be computed as a weighted sum of each value function basis. This enables instant zero-shot recomputation of the value function for any reward function which exists in the span of the reward bases. Moreover, this can be achieved using only classical TD learning methods in parallel for each reward basis.

### Reward Bases

Instead of assuming a unified total reward function *r*(**x**), we assume that the reward function is instead a linear combination of component reward functions, or *reward bases*,

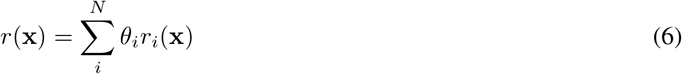

In the above equation, *r*_*i*_(**x**) denotes an individual reward basis and is an arbitrary function which assigns scalar rewards given states, *θ*_*i*_ is a scalar weighting coefficient for each reward basis, and *N* is the number of reward bases. We can then substitute this definition into our definition of the value function (Equation 2), and immediately derive that the value function also decomposes into a weighted sum of component value functions,

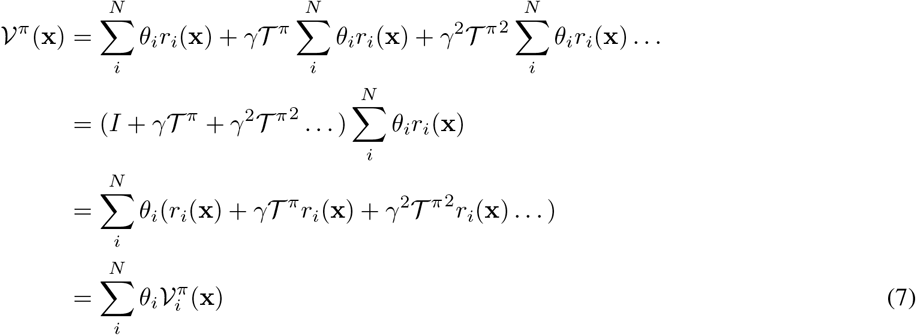

where we define 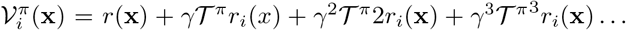 as a *value function basis* since it is the value function of one specific reward basis. All of these value function bases can be learnt using the exact same TD learning rule which is used to learn the total value function, but using the reward basis as the reward instead of the total reward. The crucial point is that all the value function bases can be learnt in parallel using a single set of state-action-reward tuples. We can estimate each value function basis using a version of the Bellman recursion for each basis, namely,

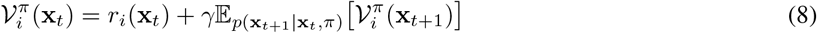

And similarly derive a TD learning rule to estimate each value function basis based on the RPE for that basis,

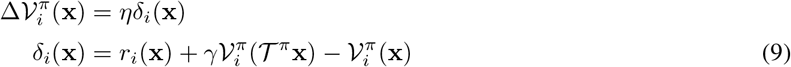

Thus, given a set of reward and value function bases, and a set of weighting coefficients {*θ*_*i*_}, we can instantly compute any value function spanned by the reward basis dynamically. What this enables, in practice, is zero-shot generalization of the optimal policy as computed from the value function for *any* reward function which can be expressed as a linear combination of the reward bases. Importantly, the reward basis functions themselves can be arbitrarily nonlinear functions of the state, so that nonlinear reward functions can still be learnt, as can nonlinear changes to the reward function. In terms of computational cost, the cost of the reward bases computation is linear in the number of bases chosen – i.e. *𝒪* (*N* · | *𝒳* |). This is simply because each reward and value basis must be computed in parallel before being combined into the final value function ^5^. Although described as weights, the weighting coefficients {*θ*_*i*_} are not necessarily synaptic weights to be learned but instead must change dynamically to match the changing physiological state. This therefore requires a control system to set these weighting coefficients. Such a control system is simply assumed in this paper, but a potential implementation based on homeostatic set-points is discussed in the Discussion section.

To build intuition for how the reward basis model functions, in Figure 1 we present a visualization of the value functions learned by a TD and reward basis agent on the room task presented later in the paper. The agents must navigate around a 6 × 6 grid where they can obtain rewards in three locations (indicated in yellow). We assume that these three rewards are of different types, and activate separate reward bases (Figure 1B top). The TD agent learned a total reward function where all three locations were equally desired composed of defined as 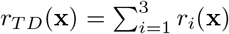 (Figure 1A top). The value function learnt by the TD and reward basis agents are shown in the bottom row of Figure 1. Using the set of value bases in Figure 1B and equal weighting coefficients of 1, the reward basis agent can exactly reconstruct the total value function of the TD agent (cf. Figure 1A bottom and Figure 1B bottom right). Importantly, the reward basis agent can also instantly generalize to other reward preferences. For instance, when only the first reward becomes valuable and the others are not, the reward basis agent can choose its actions according to 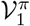. Thus starting in the bottom left corner of the environment, it would instantly go up. By contrast, the TD agent only maintains *𝒱*^*π*^, so it would go right, and by additional experience in the environment could learn that this action is no longer rewarding.

**Figure 1:**
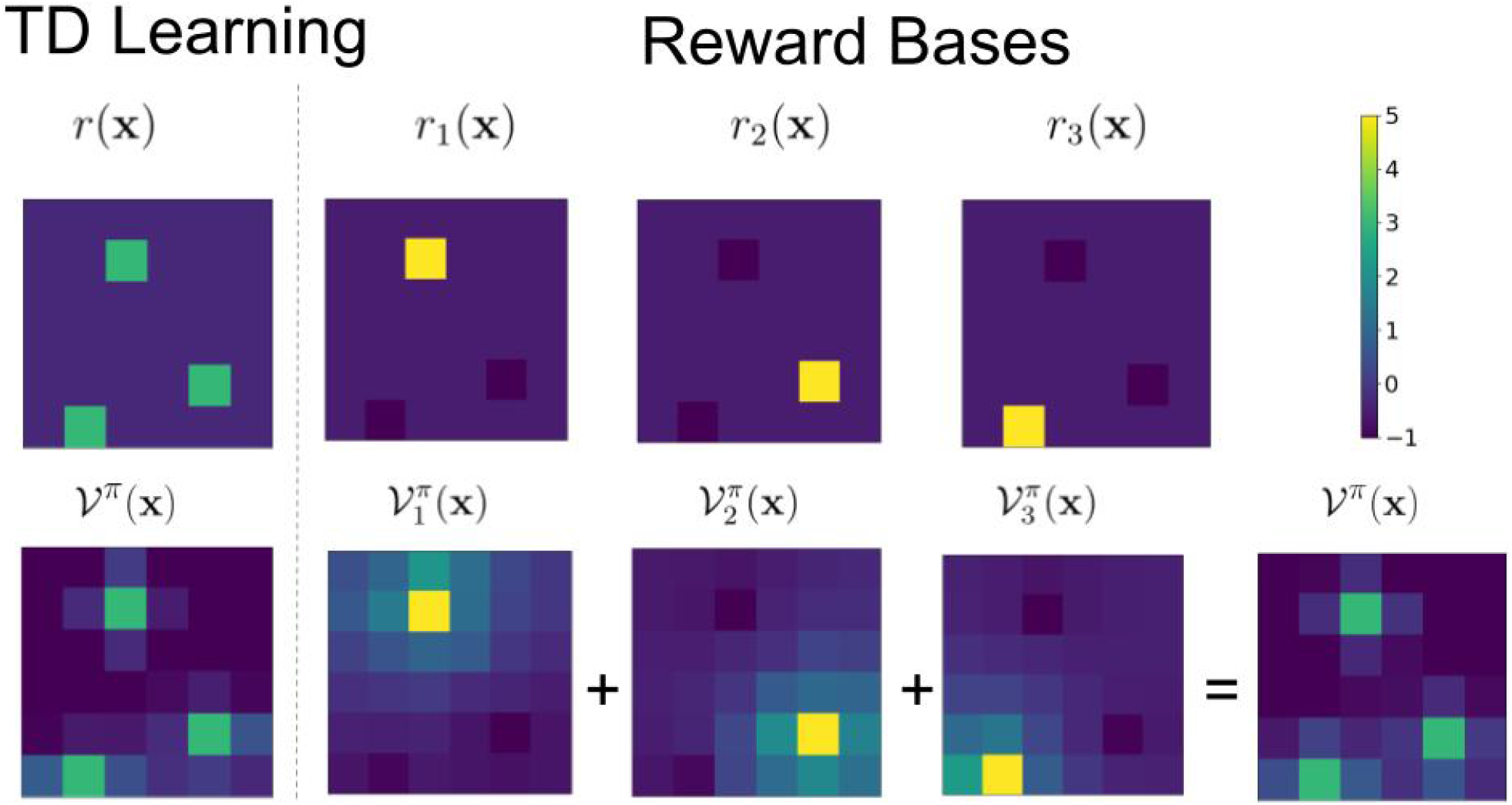
Visualization of the reward basis model using reward and value functions computed for the room task described later in this paper. The task consisted of a 6 × 6 grid with three items. The reward basis agent used a separate reward basis for each object, giving it +5 for that object, −1 for the other objects and −0.1 for all other squares. For the TD agent, the total reward was equal to the sum of the three reward bases. The value functions displayed are obtained by the agents exploring the environment for 1000 steps with a random action policy.

#### Neural Implementation with State-modulated Prediction Errors

In the model described in the previous section, the reward bases *r*_*i*_ and the weighting coefficients *θ*_*i*_ are decoupled. The rewards and reward prediction errors are still computed and the value basis is still updated even if that basis is not valued at the current moment. This is mathematically optimal given no computational or resource constraints since it enables flexible and instant generalizations to the greatest possible range of reward revaluations. However, there is a growing body of research that suggests that this is not the case in the brain since animals only appear to learn value functions of tasks when they are important to the animal or have motivational salience. For instance Aw, Holbrook, de Perera, and Kacelnik (2009) show that when fish are first exposed to a stimulus when hungry, they will chose it substantially more often over an equally palatable alternative than when they were exposed to the stimulus when sated. This implies that there is a direct modulation of the values learnt during training based on the current physiological state which is maintained even after the physiological state has changed. Cone et al. (2016) has similarly shown that physiological state can modulate dopaminergic prediction error responses such that when the animals are trained to receive food rewards after a cue in a ‘depleted state’ i.e. they are hungry – in testing, dopaminergic neurons respond to the cue (CS) and not the food reward (US) as in the classic results of (Schultz et al., 1997). However, when the animals are trained in a sated state, when tested the dopamine neurons respond only to the food reward (US), ‘as if’ they had not learned the task at all.

The version of reward basis model described so far cannot explain these findings since it decouples dopaminergic activity – and hence value function learning – from the physiological state such that the state is only used to dynamically modulate the weighting of the value function bases during action evaluation. However, as shown by van Swieten and Bogacz (2020) such effects can be straightforwardly explained by redefining the dopaminergic prediction errors such that they are also modulated by the physiological state. Effectively, this means that the ‘learning rate’ of the TD update is modulated by the physiological state variable such that animals learn much more rapidly when they are ‘depleted’ and much less rapidly when they are sated. van Swieten and Bogacz (2020) demonstrated that such state-modulated prediction error can also be viewed as error in prediction of subjective utility of rewards given specific mathematical assumptions about the form of the utility function. These insights can be straightforwardly incorporated into the reward basis model by adding motivational salience to the prediction errors directly over and above the modulation of the value bases, which greatly aids its neural plausibility. Mathematically, this model can be expressed by defining a ‘modulated prediction error’ 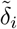,

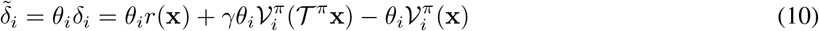

The TD update can then be derived as a gradient descent on the squared prediction errors,

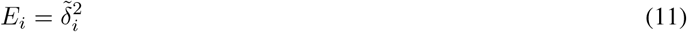

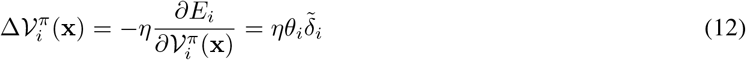

This results in a modified TD learning rule for the value bases which effectively defines an adaptive learning rate schedule where the learning rate depends on the physiological state. Although slightly complicating the algorithm, this approach has several potential advantages for the brain. Firstly, it provides a natural scaling of the learning rate with physiological stress, so that the learning rate is higher when the physiological state is more perturbed and hence the *θ*_*i*_ weightings are higher. This has clear advantages since it is important to learn fast in such cases. On the other hand, having a reduced (or no) learning rate in the case of satiation may also be beneficial. Reducing the learning rate may reduce the metabolic cost of making the updates, since less synaptic plasticity is required and the brain has been heavily optimized by evolution to minimize energy expenditure (Sterling & Laughlin, 2015). Secondly, this modulation can provide a natural pruning mechanism for the brain to learn to ignore (by setting *θ*_*i*_ = 0) physiological dimensions that are rarely important to the animal. This pruning may allow the animal to learn over time a set of reward bases that can accurately track common changes in the reward function but without representing a large number of redundant or extremely rarely used reward bases. In this paper, we use this modified learning rule to match the dopamine responses observed in the ventral striatum.

This state-modulated prediction error model also possesses a potentially straightforward implementation in the circuitry of the basal ganglia. This is achieved as a simple modification of the original circuit model of Schultz (1998). In this model, dopaminergic neurons in the VTA receive two inputs – the incoming reward and the temporal derivative of the value function. To compute the RPE, the dopaminergic neurons simply need to subtract these two inputs. Figure 2A represents this model. These RPEs can then be sent to the ventral striatum where they are used to update synaptic weights encoding the value function. It is instructive to first consider how the computational version of the reward basis model without state modulated prediction errors could be mapped on a neural circuit - although we pointed that it is unlikely this model describes vertebrate basal ganglia, it may still capture reinforcement learning in simpler animals (see Discussion). Moreover, this architecture is of independent computational interest for applications in machine learning. The mapping of the algorithm on a neural circuit is shown in Figure 2B, and the only change is that instead of assuming a single homogenous population of dopamine neurons responding to a global reward signal, we instead assume that neurons encoding values and prediction errors are parcellated into groups which each respond to a single reward basis. This reward basis parcel is simply the same circuit as original model in miniature in that dopamine neurons simply receive both the reward for that basis, and the temporal derivative of the value function for that basis and computes the RPE for that basis, which is then sent separately to the regions which require it. A set of parallel units represents the value function bases, from which the ultimate value function can be computed through a simple linear combination with the weighting coefficients. However, in this model the weighting coefficients do not affect dopaminergic activity as they appear to do in the vertebrate brain. Finally, in the state-modulated prediction error model (Figure 2C), the weighting coefficients directly modulate the dopaminergic responses in the VTA as well as striatal activity, thus defining, in effect, an adaptive learning rate for the value function updates.

**Figure 2:**
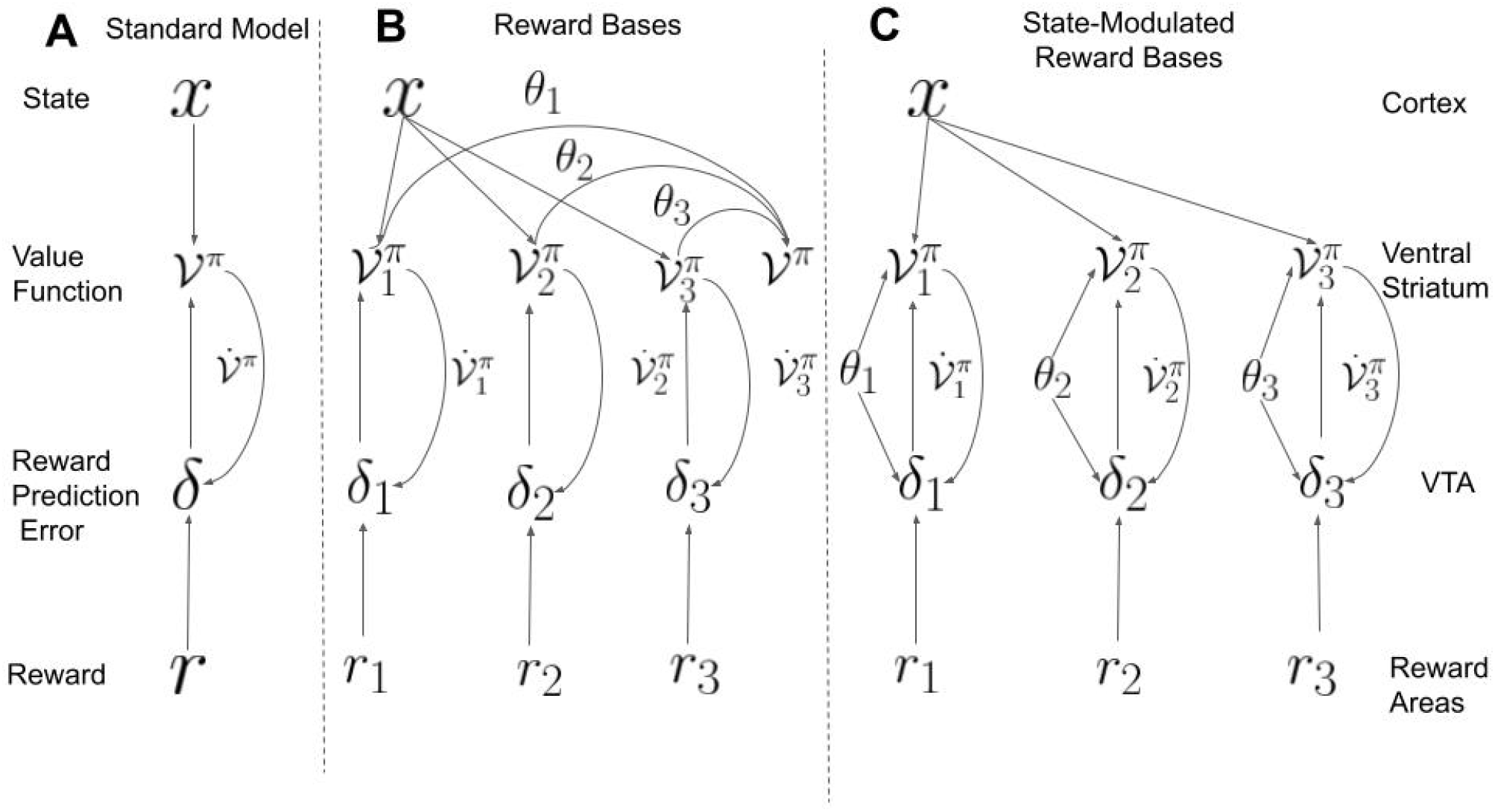
Comparison of standard theory with the reward basis model and the state-modulated reward basis model. **A**: Schematic Representation of the standard RPE/TD learning of dopamine and striatal function. Dopaminergic neurons in the VTA receive reward input and compute a RPE. This RPE is projected to the striatum which implements a TD update like rule at the corticostriatal synapses the learn a value function. **B**: Schematic of the reward basis model. This requires parallel neural populations of computing prediction errors and ‘value function’ neurons with the same connectivity patterns as the standard model, in additional to a further summation step where the value function bases are summed together with the weighting coefficients. **C**: Schematic of the state-modulated reward basis model. Instead of weighting occurring during output summation, the homeostatic state values directly modulate the activity of the striatal and dopaminergic neurons. This results in learning being sensitive to homeostatic state at the cost of being unable to learn about value bases which have 0 weight. Dopaminergic VTA neurons receive reward information from many distinct brain regions (Tian et al., 2016) since there are many distinct types of reward and rewarding stimuli that must be collated and compared.

As explained in detail in Discussion, the reward basis model makes several clear experimental predictions. The fundamental prediction is that, under the assumption that dopaminergic neurons primarily compute RPE, it would predict that individual dopamine neurons signal RPEs for a specific reward basis. Secondly, given the ubiquity of topographically organized representations in the brain, the dopaminergic neurons representing specific reward bases may be spatially clustered. We will evaluate the evidence for separate populations of dopaminergic neurons, as well as further experimental predictions made by the model in the Discussion section.

### Reproducing Experimental Data

#### Dead Sea Salt Experiment

A classic experiment that demonstrated zero-shot reward revaluation abilities was conducted by Robinson and Berridge (2013). They utilized a Pavlovian association paradigm in which rats were repeatedly exposed to one of two cues – associated with either intra-oral delivery of sucrose (pleasing) or salt (aversive) solution. The rats were trained in a food-restricted state but when injected with aldosterone and furosemide which mimics severe salt deprivation, immediately responded positively to the salt cues. These results imply that the rats clearly possess the ability to perform instant (zero-shot) generalization to update associations upon physiological change with no direct experience of the positive rewards associated with the change. Such an phenomenon cannot be explained by TD models of reinforcement learning, since the animals never experience the salt solution as rewarding. This means that there are no prediction errors to drive updates to the value function. These results can, however, be directly explained by the reward basis model.

In our model, since salt is a physiologically important quantity (Krause & Sakai, 2007), we assume the animals would maintain a ‘hardwired’ reward basis recording salt prediction errors and computing a salt value function during the Pavlovian association phase. However, upon the injection of the aldosterone and furosemide, the reward basis model can dynamically modulate its (negative) salt value function with its physiological state to perform instant revaluation of its associations with the salt lever. This can occur even in the absence of any positive reward signal obtained by experiencing the salt, since the weighting coefficients *θ*_*i*_ act directly on the value function bases, and thus the value bases themselves do not have to be updated. We demonstrate this zero-shot generalization capability of the reward basis model by replicating the paradigm of Robinson and Berridge (2013) in simulation and matching the experimental behavioural results obtained.

In simulation, the rats were exposed to interactions with the salt or juice levers (at random) and learnt a value function of the salt or juice. We assumed a linear proportional relationship between the learnt value function and degree of appetitive behaviour towards the lever. Figure 3 shows that in the homeostatic (normal) condition, both temporal difference (TD) learning and reward basis learning are able to reproduce the behaviour of the animal, while in the sodium depletion condition, the reward basis agent can instantly generalize to match the behaviour of the agent, thanks to the weighting coefficients *θ* while the TD agent has no mechanism to vary its value function according to its physiological state, and so simply predicts that the salt lever will remain aversive to the animal in the sodium depleted condition.

**Figure 3:**
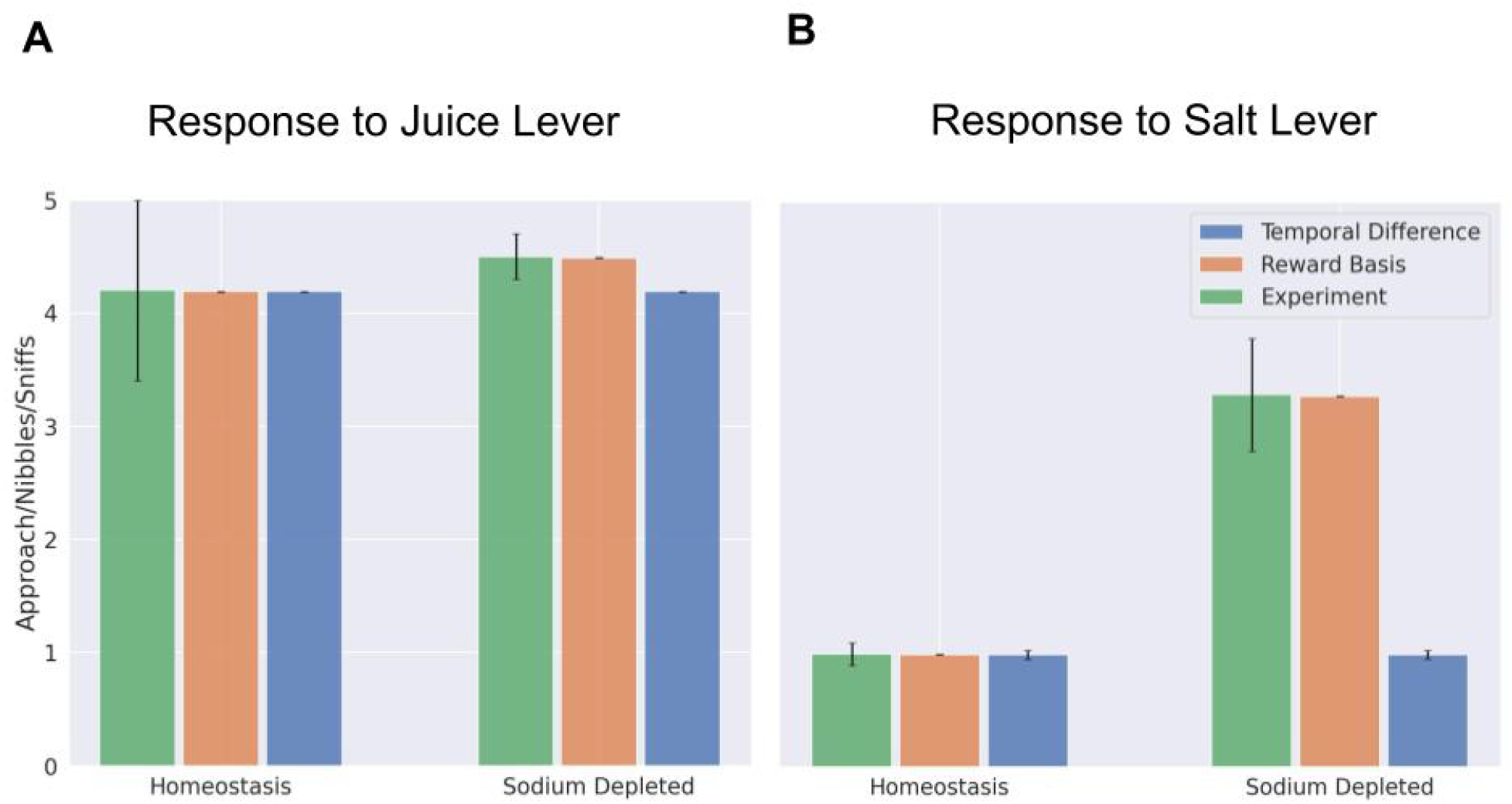
Simulated replication of results of Robinson and Berridge (2013). **A**: Approaches / appetitive actions towards the juice lever after conditioning in the homeostatic (normal) and salt-deprived conditions. **B**: Approaches / appetitive actions towards the salt lever after conditioning in the homeostatic (normal) and salt-deprived conditions. Experimental data is replotted from Figure 3C of (Robinson & Berridge, 2013).

For additional understanding of the performance of the reward basis model against TD learning in similar reward revaluation tasks, we also simulated a variant of the Robinson and Berridge (2013) experiment as a two-armed bandit task in which the animal could choose to go to a lever and either receive juice or salt. In this scenario, we can directly simulate the actions of the animals after reward revaluation in extinction (no reward provided after revaluation) or when reward is provided. We see that in all cases the reward basis algorithm provides instant generalization while TD learning requires a number of interactions with the environment to relearn the value function given the new reward function. For these experiments, please see Appendix A.

#### Predicting Dopamine Responses

Another key criterion of the biological plausibility of our model is whether it matches observed data on dopaminergic signalling in the brain. Here, we focus on qualitatively matching the results of Papageorgiou, Baudonnat, Cucca, and Walton (2016) who used fast-scan cyclic voltammetry (FCV) to measure dopamine levels in the Nucleus Accumbens Core (NAcc), part of the ventral striatum, in a reward devaluation task in which rats chose between options associated with either a food or a sucrose reward under varying conditions of selective satiation. FCV measures relative changes in extrasynaptic dopamine. Thus, we use our model to make predictions of dopamine release in the NAcc, which we take to be the sum of the dopaminergic neurons, i.e, 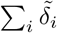, summing across the contributions of all reward bases.

A schematic of the experimental paradigm of Papageorgiou et al. (2016) can be seen in Figure 4A. The experimental data were taken from forced-choice trials on which the animal always knows at cue onset which choice is available. To model these data, we simulated the choices and rewards of an agent interacting with the Papageorgiou et al. (2016) paradigm illustrated in Figure 4A where the animal could choose either a food or a sucrose lever which usually gave out the main reward type (0.8 probability), and either gave MORE food (4 units instead of 1) with 0.05 probability or switched to 1 unit of the other reward type with 0.15 probability. The reward basis agent was trained with two reward bases, one which gave out 1 reward for each food reward and 0 for sucrose, and a sucrose basis with the opposite reward schedule. The value functions for each reward basis were trained under a random policy. After training, the animals also were tested in devaluation sessions where they were fed to satiety in one reward type but deprived of the other. In these trials the reward basis weights *θ*_*i*_ were fitted to the data using a maximum likelihood scheme to determine how the reward weights changed during devaluation.

**Figure 4:**
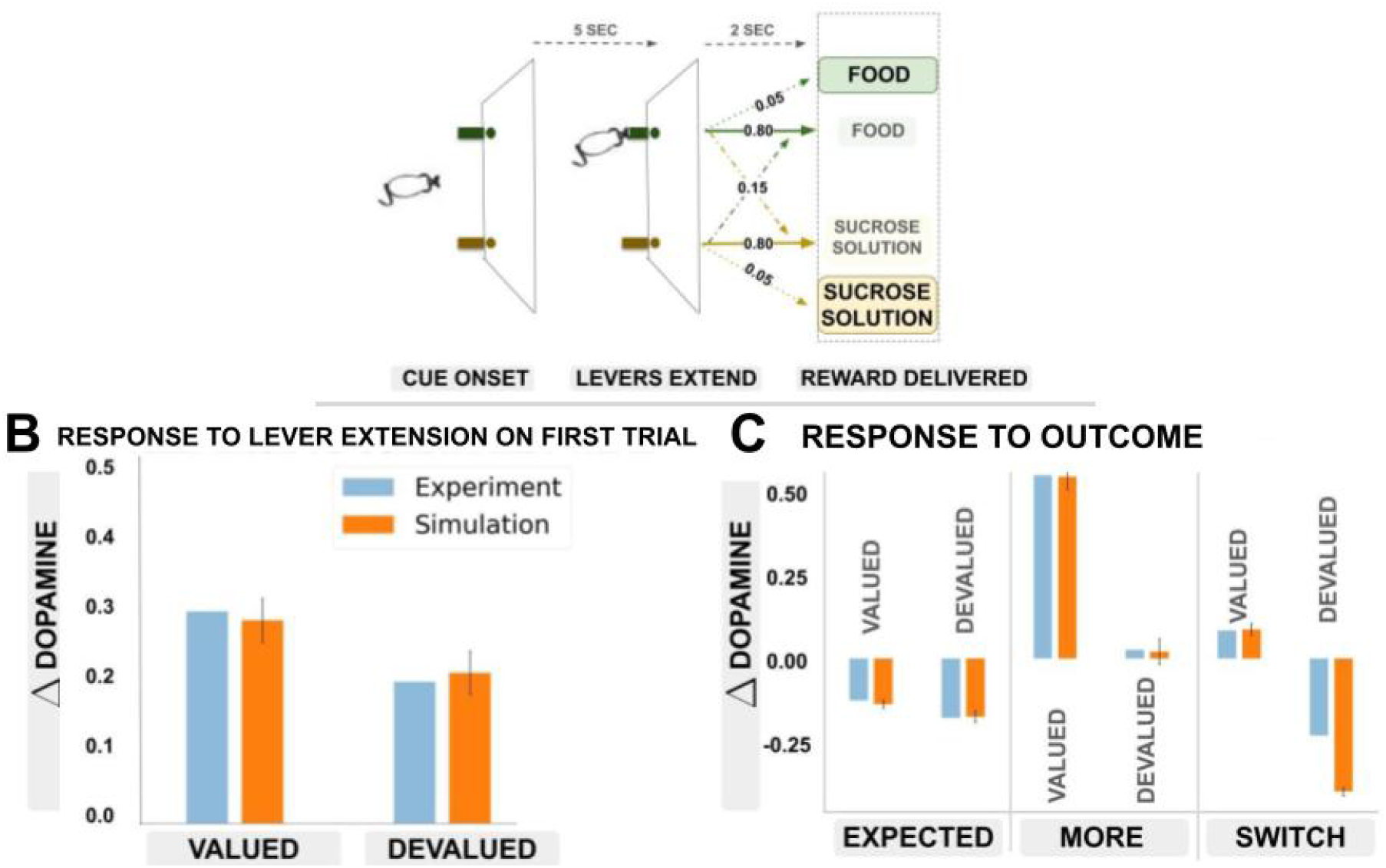
Dependence of dopaminergic responses on physiological state. **A**: The experimental paradigm of (Papageorgiou et al., 2016) (adapted from original figure). Animals were able to choose between two different reward types – food and sucrose – which could be obtained at two different levers. In free-choice trials both rewards were available, however in forced-choice trials, only one of the levers delivered its reward type. At the onset of each (forced) trial, animals were presented with a cue indicating which of the two rewards they were able to receive. Animals were required to go to the correct poke and stay for a set length of time before the reward was delivered. In some conditions, the poke did not deliver the expected reward but either four times the amount of reward (MORE condition), or the other reward type (SWITCH) condition. In the devaluation condition, the animals were fed to satiation in one of the reward types but deprived of the other. **B**: Comparison of simulated results against data taken on the *first trial* just prior to reward delivery. These results show that devaluation can affect dopamine levels prior to any experience with the devalued reward, as also found in (Robinson & Berridge, 2013). **C**: Comparison of simulated results against (Papageorgiou et al., 2016) in the MORE and SWITCH conditions. Experimental data levels computed as the levels at the end of the cue or reward delivery period. Dopamine levels are benchmarked relative to dopamine at cue onset. Simulation data comes from a simulation of the experimental paradigm and error bars are the standard deviations over 20 runs.

In Figure 4B, we first study the dopamine response levels at cue period relative to cue onset on the *first trial* after devaluation before the animals have any experience with the devalued reward. There are reliable differences in dopaminergic activity following presentation and choice of the devalued option relative to the valued one even on the first trial, a finding which cannot be accounted for by standard TD learning methods. Our method, however, straightforwardly predicts these results. This is because the weighting coefficients are lower for the devalued case, thus the total dopamine level will be decreased when the devalued stimulus is cued even before the devalued outcome is delivered since the positive prediction error induced by obtaining the devalued reward will be outweighed by the larger negative one induced by not obtaining the valued reward.

In Figure 4C we next compare the experimental to simulated responses to outcome in three conditions. The first condition is the level of dopamine when the expected reward type is presented for both valued and devalued rewards. The experimental data shows that the devalued option elicits a slightly lower level of dopamine than the valued option. We see that our model can accurately reproduce the qualitative relative levels of dopamine observed between the two conditions with the dopamine being higher in the case of valued rewards than devalued. In particular, the model produces slightly negative dopamine level after receiving valued reward. This is because, the simulated dopamine level is a sum of RPEs for two bases: receiving the expected amount of valued reward causes a zero prediction error for the valued reward basis 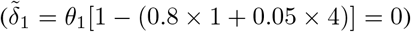, and a negative RPE from the devalued reward basis 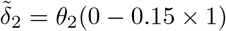 which arises from the SWITCH trials. Conversely, receiving the devalued reward elicits a lower level of dopamine activity in the striatum because of higher weighting of the negative prediction error from not receiving the valued reward that would be given if a trial turned out to be a SWITCH trial.

Secondly, we compare the activity of the valued and devalued stimuli in the MORE condition, where animals receive a larger amount of the expected reward type. Here, the reward basis model can accurately reproduce the qualitative relative levels of dopamine observed between the two conditions with dopamine being higher in the case of valued rewards than devalued. This is because receiving the valued reward causes a positive prediction error scaled by a high weighing coefficient, which lets it outweigh the simultaneous negative prediction error caused by not receiving the devalued reward. Conversely, receiving the devalued reward elicits a lower level of dopamine activity because the positive RPE is scaled down while the negative prediction error from not receiving the valued reward is scaled up.

Thirdly, the reward basis model reproduces the *SWITCH* condition, where the expected amount of the alternative reward type is delivered. For the valued reward, the positive prediction error caused by the switch is scaled by a large weighting coefficient. On the other hand, the model predicts a large reduction in dopamine levels for the switch to the devalued reward. Since there is a large negative prediction error with a large weighting coefficient associated with not getting the valued reward, while the positive prediction error for getting the devalued reward is scaled by a low weight, this leads to the negative prediction error outweighing the positive prediction error, resulting in a depression in dopamine. However, this effect does not appear as prominently in the experimental data, which found that the devalued switch condition elicits a response roughly equivalent to the expected devalued reward. In some intuitive respects, however, the Papageorgiou et al. (2016) result in this condition is more surprising than our model’s prediction. Specifically, our model predicts that expecting the valued stimuli and receiving the devalued stimuli instead is somehow worse (in dopaminergic terms) than expecting to receive and ultimately receiving the devalued reward in the first place. This accords with psychological intuition where having something you expect taken away from you is worse than never having expect in the first place. On the other hand, this intuition does not accord with the results from Papageorgiou et al. (2016) which suggest that the rats do not respond in such a way but treat switched-devalued rewards in the same way as expected devalued rewards.

### Comparison with alternative models

In this section we perform a computational evaluation of our model against competing models (successor representations and a homeostatic model) in a more complex task with multiple distinct reward bases. This allows us to demonstrate how the reward basis algorithm allows for natural generalization across tasks, as well as how it performs equivalently to the successor representation at a smaller computational cost, while being able to solve multi-step tasks that defeat the homeostatic model. We setup our environment to be a small grid-world room, with multiple different squares which each represent different types of objects. The goal is, when started in a random position in the room, to reach a specific type of object as fast as possible. The room consisted of a 6 × 6 grid-world and the object positions were randomly initialized at the start (Figure 5A). Agent training was separated into episodes such that whenever the agent reached the correct object, the episode would end, and the agent would restart in a randomly chosen square of the room. The value function was a 6 × 6 = 36 dimensional vector which was initialized to 0 for all agents and all states. In our experiments there were three different types of objects, which were each placed in one random square in the room. The agent could move a single square left, right, up, or down in each time-step. The room was bounded by ‘walls’ such that if the agent tried to move left when it was already at the edge, it would stay where it was instead of wrapping around. The reward basis agent used three reward bases which each gave +5 for the object they were associated with, −1 for the other objects and −0.1 for all other states. There were three reversals (changes in objects’ desirability) – making each of the three objects the ‘correct one’ while demoting the others to incorrect.

**Figure 5:**
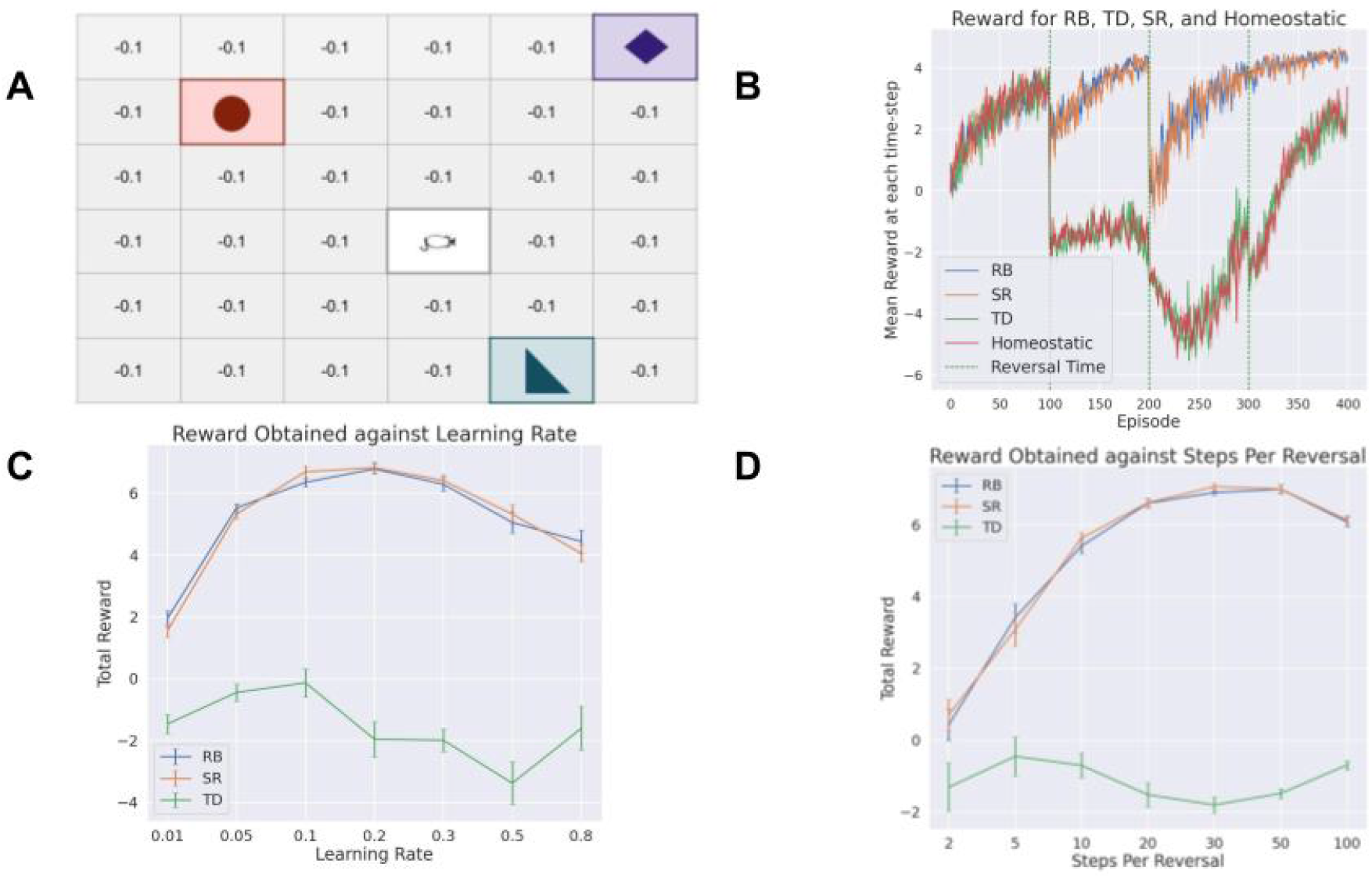
Room task. **A**: Schematic of an example instance of the room environment. The agent begins at a random point in a 6 × 6 room and must reach one of the red, green, or blue squares, receiving +10 if it reaches the correct one and -5 if it reaches an incorrect one. **B**: Comparison of the reward obtained by the temporal difference (TD) agent, the reward basis (RB) agent, the successor representation (SR), and the homeostatic agent on an example run of the room task. The vertical dashed lines represent the reversals. **C**: Performance obtained by RB, TD, and SR agent over a range of learning rates with 50 episodes between each reversal. **D**: Performance obtained by RB, TD, and SR agent over a range of trials per reversal (to investigate learning speed since to achieve high reward in fewer trials before reversal requires faster learning). A learning rate of 0.1 was used. All error bars represent standard deviation over 10 runs.

In Figure 5, we plot the performance of the reward basis agent against two competing models on the room task. First, we compared its performance against the ‘homeostatic model’ of Zhang et al. (2009). While full details can be found in the Methods section, this model aims to simulate homeostatic updating of behaviour by modulating the expected reward term with a homeostatic factor *κ*(*u*) where *u* represents the homeostatic state. Since it directly modulates predicted rewards, it can solve single-step tasks such as the dead sea-salt experiment but cannot generalize behaviour across multi-step tasks with identical rewards on a single step, since the homeostatic factor will modulate all such rewards identically. The fundamental reason for this difficulty is that it does not generalize across *value functions* which is necessary to correctly adapt long-term behaviour upon changing the reward function. An identical critique applies to the homeostatic model of Keramati and Gutkin (2014) who define the reward function as the sum of distances to various homeostatic set-points, but which also does not generalize across value functions.

Secondly, we compare our model to successor representations which can generalize correctly across changing reward functions. Successor representations (Dayan, 1993) is an alternative model-free RL method that also enables instantaneous generalization across changing reward functions. They work by learning a ‘successor matrix’ based on discounted state occupancies (see Methods section for mathematical details) which can then be combined with the reward function to yield the value function. In the limit, therefore, the reward basis algorithm and successor representations have approximately equivalent capabilities. However the reward basis algorithm has several computational advantages. Firstly, instead of storing a | *𝒳* | × | *𝒳* | successor matrix, it only requires | *𝒳* | × *N* memory where *N* ≤ | *𝒳* | is the number of reward bases. Moreover, the evaluation of the value function with the successor representation requires an *𝒪* (𝒳) summation while the reward basis agent only requires *𝒪* (*N*) complexity summation per value function evaluation.

In Figure 5B, we can see the performance of the homeostatic agent of Zhang et al. (2009). We see that, as expected, this agent performs comparably to the temporal difference agent and fails to generalize across the changing reward function in this task. This is because the homeostatic model cannot update its value function when the reward function changes and hence does not generalize.

In Figure 5B, we also plot the performance of the reward basis agent against the successor representation agent and a TD learning agent. We observe that both the reward basis agent and successor agent rapidly generalize to the reversal while the TD agent must instead gradually relearn the task each time based on multiple interactions with the environment. Both the reward basis and successor agents still also require some retraining after reversals likely due to the approximate nature of the learnt value functions or successor matrix given their limited number interactions with the environment.

In performance on this task, the reward basis and successor representation agents both achieve approximately equal performances across a wide range of parameter settings (Figure 5C-D). However, the reward basis agent is substantially computationally cheaper, especially in terms of memory cost. This difference in the required memory for the reward basis algorithm vs successor representations can be significant. For instance, in the small task presented here, with three reward bases, the number of values stored by the reward basis algorithm was 36 × 3 = 108 while the successor matrix required 36 × 36 = 1296 – a substantial increase. This advantage will generally only increase with state-size since, in general, where the number of reward basis is small, the memory cost of the reward basis algorithm scales linearly while the successor representation scales quadratically. In Appendix B, we provide further investigations of the computational properties of the reward basis algorithm compared to successor representations as well as an additional experiment demonstrating the functioning of the algorithm with mixed value bases.

It is also possible to show mathematically that the reward basis model is closely related to the successor representation and, indeed, can be intuitively thought of as a compressed successor representation with rewards tuned only to relevant dimensions. In the successor representation (see Methods section for more detail), the fundamental element is the successor matrix *M*^*π*^ which is an | *𝒳* | × | *𝒳* | matrix which stores the discounted state occupancies for a given policy. This means that *M*_*i,j*_ stores the discounted total probability of moving from state *i* to state *j* during an episode. If we possess this state occupancy matrix *M*, we can then compute the value function with a simple matrix multiplication,

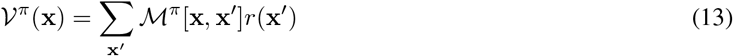

To understand how the successor representation relates to the reward basis scheme, note that each value basis can be expressed in terms of the successor matrix similar to the total value function,

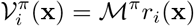

This can be demonstrated as a straightforward consequence of the linearity of the Bellman equation for a fixed policy. By the standard successor representation decomposition of the value function (see methods),

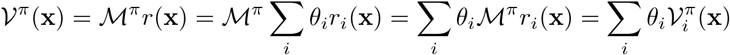

While this decomposition eschews the computational benefits of the reward basis model compared to the successor representation, it is valuable to gain intuition as to the relationship of the reward basis model with the successor representation, and helps explain why the reward basis model achieves equivalent performance to the successor representation at a significantly smaller computational cost.

## Discussion

In this paper, we have proposed a novel mechanism for zero-shot generalization to changing reward functions in reinforcement learning tasks which relies solely on model-free TD learning methods. Crucially, our results demonstrate that behavioural flexibility in the face of changing reward functions does not require model-based evaluation methods as argued in Dayan and Berridge (2014) but instead can be handled in a purely model-free manner using a straightforward extension to the classical TD learning method which is already widely believed to be implemented in the basal ganglia. Moreover, this method is extremely straightforward and can be applied to any reinforcement learning method which estimates value or Q functions. With a reasonably small number of reward bases (less than the full size of the state space), the reward basis method maintains the computational efficiency of model-free RL while also being substantially less computationally and memory expensive than successor representation methods, which must materialize an | *𝒳* | × | *𝒳* | successor matrix. We have demonstrated that our approach allows for almost flawless zero-shot reward revaluation performance in a variety of tasks, including several behavioural tasks used in neuroscience experiments such as the seminal Dead Sea salt experiment Robinson and Berridge (2013), where our approach can match the capabilities of animals to perform zero-shot reward revaluation using only temporal difference methods.

### Experimental Predictions

A key experimental prediction arising from our model is that different dopaminergic neurons in the mid-brain should be differentially sensitive to different reward types since, according to our model, they are computing RPEs for different reward bases. It is important to note that it is not necessarily the case that the reward bases actually encoded in the brain will be directly as interpretable as the examples presented in this paper such as a ‘food neuron’ and a ‘sucrose neuron’. All that is needed is that the population of reward bases suffice to span the space of reward function encountered in the environment, not that each neuron encodes a specific kind of stimuli. It is very possible that there is instead a neuron which represents 0.3*sugar* + 0.6*salt* + 0.1*water* and so on. Nevertheless, a key prediction is that given a specific stimulus, there should be populations of dopaminergic neurons which respond differentially to the same reward. Moreover, by varying reward stimuli combinations, it may be possible to actually decode the reward bases implemented in the VTA – i.e. figuring out *what linear combination* of primary rewards a reward basis neuron actually encodes.

An additional, albeit more speculative, experimental prediction is that if the dopaminergic neurons in VTA are parcellated into bases which form spatial clusters (i.e. all dopaminergic neurons for reward basis *i* are close together instead of being uniformly spread throughout VTA), then we should expect topographically organized projections both to and from these specific parcels of reward basis neurons in VTA. This is because each cluster of dopaminergic neurons likely needs different incoming information to compute its specific RPE. For instance, neurons associated with a basic homeostatic reward basis, like one identifying the presence or absence of salt primarily, need inputs from areas like the lateral hypothalamus and the gustatory cortex in the insula. However, a more complex reward bases, perhaps attuned to positive social emotions, will likely require primarily cortical inputs from prefrontal, orbitofrontal, and cingulate cortices. A clear prediction of the reward bases model is there should be topographically separate projections into the VTA with different regions of the brain projecting to spatially separated regions of the VTA, while the global RPE framework of Schultz (1998); Schultz et al. (1997) should predict the opposite – namely that all regions of the brain which project to the VTA should project uniformly throughout it so that there is no spatial structure since every RPE neuron requires precisely the same inputs. Our model would also predict parcellated and topographically separate projections *from* VTA are also possible, although this is not necessarily the case since it could be that the value function bases are computed and combined together in the same regions – for instance in the ventral striatum – although there might nevertheless be spatial structure to the projections from VTA to different regions of the ventral striatum where each value function basis is computed.

#### Relationship to Neural Data

Although it is not currently clear whether differentially responsive dopamine neurons exist in the mammal brain, there are some interesting preliminary findings which point in this direction. While initially it was thought that RPEs encoded by dopamine neurons in VTA were largely homogeneously encoding the same RPE, recent work has begun to add some nuance to these conclusions. There is also evidence for dopaminergic neurons in the tail of the striatum which encode RPEs specifically for aversive stimuli (Brischoux, Chakraborty, Brierley, & Ungless, 2009; Watabe-Uchida & Uchida, 2018) – essentially negative RPEs. These neurons appear to have different connectivity patterns than the standard dopamine neurons and thus may constitute a separate class (Verharen, Zhu, & Lammel, 2020). This has been argued to signify that the classical reinforcement learning system is actually split into two axes – a positive axis which encodes rewards, and a negative axis which encodes costs and negative quantities (Schultz, 2019; Watabe-Uchida & Uchida, 2018).

Finally, there is growing evidence for substantial heterogeneity in response properties of dopamine neurons in terms of their response properties to rewarding and non-rewarding stimuli. For instance, recent work has demonstrated convincing evidence that dopaminergic neurons vary in terms of their response strengths to positive vs negative rewards, thus implementing a distributional reward code which can be mapped to a posterior distribution over the value function at a population level – an approach known as distributional RL (Dabney et al., 2020; Dabney, Rowland, Bellemare, & Munos, 2018). Moreover, there is growing evidence that dopaminergic neurons are not just sensitive to reward contingencies but also various aspects of the state-space such as locomotion and kinematic behaviour (Barter et al., 2015; Dodson et al., 2016; Engelhard et al., 2018; Kremer, Flakowski, Rohner, & Lüscher, 2020) as well as choice behaviour (Coddington & Dudman, 2018; Parker et al., 2016) and indeed that this heterogeneity is topographically organized due to spatially organized cortical projections (Engelhard et al., 2018, 2019).

The idea of reward bases also fits naturally with the idea that in the brain there are multiple different kinds of reward arising from different sources. While, mathematically, the actual vectors making up the reward bases are largely arbitrary except insofar as the linear combination of the bases spans some desired space, in the brain, due to its functional specialisation and general modularity, it seems likely that reward bases, if they exist, inherit this structure. Firstly, it seems plausible that there are reward bases encoded in the hypothalamus which handle primary rewards or unconditioned stimuli. For instance, the lateral hypothalamus has long been known as the ‘feeding centre’ of the brain and contains nuclei handling many aspects of nutrient homeostasis, as well as experiments showing that lesions to the lateral hypothalamus abolish feeding behaviour to the extent that animals simply not eating until they starve to death. Other reward bases may be specific to certain fear, anxiety, or pain inducing stimuli, which would be computed in regions specialized for aversive behaviour such as the amygdala (de Oliveira et al., 2011; Jennings et al., 2013), habenula (Chan et al., 2017; Matsumoto & Hikosaka, 2007; Tian & Uchida, 2015), and periacqueductal grey (Walker, Wright, Jhou, & McDannald, 2020; Williams, Hassall, Trska, Holroyd, & Krigolson, 2017) (negative stimuli would be represented with a negative weighting coefficient, but a ‘positive’ reward basis under this scheme).

It is also possible that the brain has the capability to learn additional reward bases (this idea is developed further in a later section). For instance, cortical connections to VTA could, among other things, signal more abstract reward bases which correspond to more abstract social unconditioned stimuli such as the enjoyment of humour or friendly companionship as well as potentially additionally arbitrary bases which are constructed over time by the cortex to allow for its more abstract goals to be incorporated into the model-free basal ganglia framework. Such learned reward bases are almost certainly cortical in nature, and could be implemented in the brain through connectivity from cortex directly to midbrain dopaminergic regions such as VTA and SNc.

### Evidence in Drosophilia

Unlike in the basal ganglia of mammals where evidence for the reward basis model is still uncertain, there is a strong match between the neuroanatomy postulated by the reward basis model and that found in the mushroom body (MB) of fruitflies (drosophilia). The MB is a region involved in a range of capabilities including olfaction, associative memory, and classical and operant conditioning (Davis, 2005; Heisenberg, 2003; McGuire, Le, & Davis, 2001; Modi, Shuai, & Turner, 2020; N. K. Tanaka, Tanimoto, & Ito, 2008) and it has been intensively studied with the result that much of its neurophysiology and connectome is largely known (Aso et al., 2014; Li et al., 2020; Owald et al., 2015). The MB consists primarily of a large number (approximately 2200) of Kenyon Cells (KC) which project to a small number (32) of Mushroom Body Output Neurons (MBON). The KCs receive input primarily from the olfactory system (antennal lobe) (Owald & Waddell, 2015) but also visual and homeostatic state information (Li et al., 2020). The KCs project randomly onto the MBONs so that each MBON receives a large number of random connections from a set of KCs. There are also approximately 200 Dopaminergic Neurons (DAN)s which synapse onto the axons of the KCs and thus modulate the KC-MBON connetions. The DANs have been shown to be vital for learning novel cue-reward associations (Berry, Cervantes-Sandoval, Nicholas, & Davis, 2012; Dylla, Raiser, Galizia, & Szyszka, 2017; Kim, Lee, & Han, 2007) and appear to be represent similar RPEs as dopaminergic neurons in mammals (Bennett, Philippides, & Nowotny, 2021; Felsenberg, Barnstedt, Cognigni, Lin, & Waddell, 2017), which is unsurprising since basic neurotransmittor systems such as dopamine tend to be highly conserved over evolutionary history. Many DANs also receive direct feedback connections from the MBONs (Li et al., 2020) ^6^. Crucially, a large body of evidence, strongly suggests that the MBONs and DANs are specialized and separated into different zones or compartments which respond to and represent rewarding aspects of specific stimuli (Aso et al., 2014; Li et al., 2020; Otto et al., 2020; Owald & Waddell, 2015; Vogt et al., 2014). Specifically, there are 21 subtypes of MBONs (out of 32) neurons and each type appears to define a specific compartment while there are DANs associated with that compartment which only project to their associated MBON, and that each MBON also sends direct feedback back to that specific DAN. On the other hand, KC input to the MBONs appears relatively random across zones, although with some specialization for atypical MBONs which receive primarily visual (instead of olfactory) input.

Moreover, experimentally it has been shown that these specific DANs respond to specific reward types such as sucrose (Huetteroth et al., 2015; Yamagata et al., 2015) water (Lin et al., 2014; Senapati et al., 2019), and courtship (Cheriyamkunnel et al., 2021; Keleman et al., 2012) and appear to be instrumental in learning associations based on these specific stimuli while projecting to the specific MBON in their own compartments which could implement the value function for that specific stimulus. Crucially, if we associate the state representation **x** with the KCs, the RPEs *δ*_*i*_ with the DANs and the Value basis functions *𝒱*_*i*_ with the MBONs, then the circuitry in the MB appears to almost precisely fit a direct implementation of the abstract reward basis model previously presented, with neuronanatomically separated ‘compartments’ for a number of parallel value functions which are then combined downstream to enable flexible behavioural adaptation to changing reward functions or homeostatic drives.

Interestingly the macro-scale distinction between positive and negative valenced coding dopaminergic neurons in the vertebrate striatum is also paralleled in the mushroom body of the fruitfly, where there is a clear anatomical distinction between a cluster dopaminergic neurons in the Positive Allosteric Modulator (PAM) which generally encode positively valenced stimuli and an anatomically separate population in the protocerebral posterior lateral (PPL) which code for aversive stimuli (Aso et al., 2012; Owald & Waddell, 2015; Riemensperger, Völler, Stock, Buchner, & Fiala, 2005). Within these clusters there is further specialisation for specific positive or negative reinforcers such as food, water, or aversive odors and unpleasant temperatures. It is possible that a similar organization is preserved in mammals with two large-scale regions of positive and negative valenced dopaminergic neurons (Watabe-Uchida & Uchida, 2018) and within these regions further specialized clusters of dopaminergic neurons for specific stimuli.

Thus, there appears to be strong evidence supporting the reward basis model in drosophilia. Given that the MB is a evolutionarily highly optimized and central region of the fruit-fly brain involved in many of its more complex cognitive tasks, combined with its potential functional and anatomical similarities with regions of the vertebrate brain such as the cerebellum (Farris, 2011; Li et al., 2020) and striatum ((Waddell, 2016; Watabe-Uchida & Uchida, 2018), it is likely that its basic mode of operation has been largely evolutionarily conserved in vertebrates. This provides at least some indirect evidence for a similar mechanism, albeit significantly scaled up, being implemented in the basal ganglia of mammals.

#### Extensions to the Reward Basis model in Drosophilia

While the abstract schematic of the reward basis model provides a remarkably good match to the neuroanatomy of the mushroom body in drosophilia, and can clearly implement the fundamental computations required by the reward basis model, there are still outstanding questions regarding the representational code utilized by the Kenyon cells, the exact specifics of the synaptic updates between the DANs and MBONs, and whether that synaptic update is compatible with the standard TD update rule. An interesting additional question arises from the fact that while the MBONs appear to receive a mostly random set of inputs from the KCs, there is nevertheless some evidence of specialization of inputs, especially of the atypical MBONs which receive input primarily from visually driven KCs and appear to maintain their own somewhat separate visual channel in the MB – a motif also likely preserved in the striatum with its many parallel loops (Alexander, DeLong, & Strick, 1986). This means that, unlike the reward basis model, each value function basis is not learning from the same state information but instead sees its own individual snapshot of the state. Although potentially a problem theoretically, it is unclear whether this poses a major problem in practice, although in theory it could prevent the fly from learning arbitrary associations between different kinds of sensory information and arbitrary reward bases or internal homeostatic states.

A close analysis of the neuronatomy also reveals additional aspects that are not directly explained by our model and may, in effect, add additional levels of sophistication to the basic reward basis algorithm. For instance, a number of MBONs, as well as projecting back to the DANs of their own compartment also innervate the dendrites of DANs in other compartments, thus potentially providing a kind of reward-crosstalk or a mechanism for MBONs to modulate learning of MBONs in other compartments and it is unclear how this would affect the functioning of the reward basis model, or what the advantage of such modulation would be. Additionally, a number of specialized MBONs are known to project directly to other MBONs (Aso et al., 2014; Li et al., 2020), thus providing a recurrent link in the circuit and potentially implementing a multi-layer MBON network which thus may potentially implement more complex computations than the linear sum implemented in the reward basis model. Exploring whether replacing the linear sum with a multilayer network of value function estimates provides a computational or behavioural advantage is an interesting avenue for future work.

Finally, our proposal that the MB is fundamentally performing reinforcement learning through the reward basis algorithm differs slightly from a common perspective in the literature that the MB is involved in memory formation and associative learning (Felsenberg, 2021; Jacob et al., 2021b; Weiss & Brown, 1974). The apparent involvement of the MB in associative learning and formation since in Pavlovian association tasks learning value functions of cues is essentially identical to learning long term associative memories between cues and rewards. On the other hand, there are also examples of the behaviour of the MB which fall more towards pure associative memory and learning which do not involve rewards and are therefore more challenging to explain from a reinforcement learning perspective. It is thus possible that the MB may possess other modes of computation that allow it to implement these additional capabilities.

#### Opponent Coding of Positive and Negative Rewards

Within the fruitfly, positive and negative stimuli (inducing approach or avoidance) appear to be coded in an opponent manner where separate value functions are learnt for positive and negative associations with the same stimulus (Li et al., 2020; Owald & Waddell, 2015). Evidence for this can be seen in associative memory experiments where flies are first paired with a stimulus in a negative and then in a positive context (or vice versa) and it can be observed that two separate memories appear to be formed using different MBONs which cancel each other out rather than a single MBON adjusting (Jacob et al., 2021a, 2021b). Although speculative, this kind of opponent coding may also exist at a much larger scale in the basal ganglia system of mammals with potentially both separate populations of positive and negative dopaminergic neurons (Watabe-Uchida & Uchida, 2018). The advantages of such an opponency system as opposed to learning just a single value function include potentially more flexibility in nonstationary environments to learn different contexts for avoidance or approach behaviour as well as handling asymmetrical costs vs benefits of an action given a context. Moreover, an opponent coding model of value function learning in the context of direct and indirect pathways in the basal ganglia (Mikhael & Bogacz, 2016) has shown that such a coding allows both the mean value function as well as its mean absolute deviation to be represented simulataneously as the difference and sum respectively of the positive and negative opponent units. This thus provides an additional statistical advantage for such a coding scheme.

### Comparison with alternative models

As a comparable model-free method for instantly generalizing to different reward functions, it is worthwhile doing a detailed side-by-side comparison of the properties of the successor representation compared to the reward bases method. The advantage of the successor representation is that zero-shot reward revaluation can be achieved for *any* reward function. The reward basis method requires that the reward function be expressable as a linear combination of reward basis vectors (which themselves can be nonlinear functions of the state). Secondly, it assumes that these reward basis vectors can be specified beforehand, ideally at the beginning of the task. This means that successor methods alone can handle tasks where the reward function can change in truly arbitrary and unexpected ways over the course of a task. Reward basis models, on the other hand, work better where the space of potential reward function changes occurs in a relatively small linear subspace of the total reward space which can thus be represented well with smaller set of basis vectors than the full dimensionality of the state-space. While thus being slightly less generalizable in theory, for biological organisms, these conditions may well hold in practice such that while the reward function changes often, it typically only changes over a relatively well-known range such that a relatively small set of reward bases (relative to the total dimensionality of the world) suffice to cover the reward revaluations that actually occur. For instance, an animal may know that its reward function often changes as a function of its level of satiety, or its level of tiredness and thus these can form natural reward bases. Moreover, even if the reward function does not change solely over a linear subspace, the problem of approximating a complex reward function manifold with a set of reward bases is a simple least squares problem which can be directly solved to obtain a reasonable approximation.

On a computational level, the reward basis method is generally cheaper than the direct approach of storing the successor matrix without any approximations. This is because the reward basis method only has to store *N* basis vectors where *N* ≤ | *𝒳* | such that the number of bases is less than the total dimensionality of the state-space. This is because the number of states for an animal corresponds to the number of locations it can visit and their appearance, which is much greater than the number of physiological needs. Note that if the number of reward bases equals the dimension of the state-space, then the reward basis method can represent any arbitrary (discrete) reward function and thus has the same expressivity as the successor representation with the same computational cost. Another important advantage of the reward basis approach over successor representations is that in continuous state environments, where technically the dimensionality of the state space is *infinite*, the reward basis method is unaffected while explicitly storing the successor matrix becomes impossible and instead additional assumptions and approximations must be employed to make the method practicable (Barreto et al., 2016; Kulkarni, Saeedi, Gautam, & Gershman, 2016). Because of this, our reward basis approach generalizes much more naturally to continuous spaces and the use of function approximators such as deep neural networks since it appears that it can be immediately integrated with existing techniques for value function estimation by simply using a deep neural network to parametrize each value function basis and then linearly combining their outputs to produce the true estimated value function, whereas successor methods require a bespoke set of techniques which is still under active research (Borsa et al., 2018; Janner, Mordatch, & Levine, 2020; Machado et al., 2017) to be scaled up in this fashion.

A final advantage of the reward basis scheme over successor representations is that the evaluation of the value function in the reward basis model is ‘local’, in the sense that it only needs information about the value of the reward function at the current state. The successor representation, on the other hand, requires information about the reward function at *every other point* of the state-space to evaluate its value function. To see this, recall that the total value function for the reward basis is evaluated as 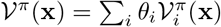 where the sum is over the value bases. On the other hand, for the successor representation the value function is evaluated as *𝒱*^*π*^ (**x**) = Σ_**x**_*′ ℳ^π^* (**x, x***′*)*r*(**x***′*) which is a sum over all potential future states **x***′* and the reward for those states, which is both a much larger sum (for *N* ≤ | *𝒳* |), and also requires the brain to be able to make reward evaluations of hypothetical states which are not being directly experienced. By contrast, the reward basis method only requires the reward function to be evaluated at the actually experienced state.

A subtle point about both the reward basis and successor representation methods is that the value functions that are learnt are only defined for a specific policy while changing the reward function naturally also changes the optimal policy, and this level of zero-shot generalization is *not* covered by the reward bases. Effectively, the reward basis value function will predict the optimal value function for the given reward given the current policy, which is not necessarily the same as the value function for a given reward function given the optimal policy for that reward function. This can become especially problematic in the case of insufficient exploration. If a region of the state-space which receives high reward under one reward basis, but is not given any reward by the current reward function and so is not explored, then switching to a reward function that heavily weights this basis will not immediately lead to a policy which prioritises the now newly-valuable regions of the state-space, since it has never explored there, and the algorithm does not know they are valuable according to this basis. While never having explored a region is obviously an extreme case, a problem also arises in the case where a region is not explored sufficiently. The value functions derived from the Bellman equation are only estimates, and if there is relatively little experience, the estimates are poor, meaning that the generalization given by changing the weighting of the reward bases will also be poor. It is important to note that the generalization afforded by successor representations also suffers from this problem, although a model-based planner with the ability to query the ‘true’ reward function in hypothetical states and a sufficiently accurate world model does not.

It is also important to clarify the distinction between the reward basis model and distributional reinforcement learning (Dabney et al., 2020, 2018). In distributional RL, the heterogeneity of dopamine neurons encodes quantiles of the RPE signal, and thus the population encodes a non-parametric representation of the posterior distribution over the value function for a single scalar reward function. By contrast, the reward basis model predicts heterogeneity in dopamine neurons in another orthogonal dimension – that of reward type – and constructs a total value function by a linear combination of value functions for each reward basis. It is also possible to straightforwardly combine distributional RL and the reward basis algorithm by estimating a distributional value estimate for each value basis in parallel.

Finally, another possibility for extending model-free TD learning methods to handle varying reward functions is to simply extend the state-space with the additional homeostatic variables. Mathematically, this would involve using model-free reinforcement learning on an extended state-space such as [x …, food_state, water_state, …] and then performing TD learning as normal. This method, however, suffers from two substantial disadvantages compared to the reward basis approach described in this paper. Firstly, it entails a substantial increase in the ultimate size of the state-space which hampers the generalizability and sample-efficiency of the resulting algorithm, and is especially acute in the presence of continuously-valued homeostatic variables which would require the animal to essentially learn its value functions from scratch for every possible setting of the homeostatic variable. Continuously varying homeostatic variables are handled naturally, however, in the reward basis scheme by just using the weighting coefficients. Secondly, this approach of extending the state-space fails to achieve the zero-shot generalization that the reward-basis scheme is capable of. This is because it uses standard TD learning which can only associate outcomes with a novel homeostatic state by directly experiencing those outcomes. For instance, in the Robinson and Berridge (2013) salt-deprivation experiments, the augmented state method would not exhibit instant generalization but would have to experience several positive pairings of the salt with the salt-deprived state to allow for this generalization – while in the original experiment the rats were specifically never previously salt-deprived in their lives to prevent this possible association developing outside the experimental paradigm.

### Learning Reward Bases

In this paper, we have just assumed a given set of reward bases which can correctly reconstruct the reward function, but the real brain faces a harder problem – that of ultimately *learning* a good set of reward bases to be able to generalize across the kinds of reward revaluations that happen often in its environment. We expect that the brain likely addresses this problem in multiple ways. Firstly, it is likely that there are a set of ‘hardcoded’ reward bases originating in the hypothalamus and midbrain which directly code for primary rewards (such as salt, sugar, water, sex etc.) as well as other survival-relevant quantities such as energetic/metabolic cost. Secondly, there is likely also a system (or multiple systems) which can flexibly *learn* new reward bases based on the kind of reward revaluations observed in the environment over the animals lifetime. Mathematically, while this is somewhat challenging, there exist methods that can achieve this (the problem is equivalent to that of learning nonlinear basis functions for regression), although precisely how they might be implemented in the brain is currently unknown. Reward basis learning, however, could be the mechanism behind phenomena such as the development of addiction to novel (evolutionarily) substances such as modern drugs. Furthermore, if and when the reward basis system fails, the brain likely also has backup model-based, as well as potentially successor-representation-based, reinforcement learning systems to allow for rapid generalization to novel reward functions at the expense of greater computational cost. An additional neuroscientific prediction that such a system would make is that there are some cues and reward function revaluations for which the animals will fail to exhibit instant behavioural generalization. This is because with a limited number of reward bases, the basis vectors cannot span the whole set of possible reward function changes. By testing empirically what sort of reward function changes can elicit zero shot generalization, as in Robinson and Berridge (2013), and which cannot, it would be possible to experimentally map the ‘reward basis manifold’ covered by the reward bases and precisely measure the extent to which the animals can generalize.

### Integration with Deep Reinforcement Learning

An additional potential strand of future work is to investigate merging the reward basis concept with recent advances in deep reinforcement learning to demonstrate zero-shot reward revaluation behaviour on challenging discrete and continuous action benchmark tasks. Another interesting direction to investigate is how this reward basis approach can be integrated with distributional reinforcement learning (Dabney et al., 2020, 2018) whereby not just the *average* of the reward estimated from a given state but instead the whole distribution or its spread (Mikhael & Bogacz, 2016). Moreover, this reward basis approach can be easily integrated with almost any reinforcement learning algorithm which computes and estimates value functions using the Bellman recursion. Specifically, all that is necessary is to run *N* value function estimates in parallel – one for each reward basis – and then to get the final estimated value function, form a linear combination of the value function bases using the weighting coefficients {*θ*_*i*_}. This means that, in theory, the reward basis approach can be straightforwardly combined with nonlinear value function estimators such as deep neural networks to implement Deep Q learning (Mnih et al., 2013), or deep actor-critic architectures (Konda & Tsitsiklis, 2000; Mnih et al., 2016) by simply maintaining a parallel set of value networks to estimate the value functions for each basis, and then computing the ultimate value function by summing across the outputs of the value function estimates with the weighting coefficients.

### Computation of the Weighting Coefficients

In this paper, we have simply assumed that the weighting coefficients {*θ*_*i*_} are known to the agent and instantly applied. This is equivalent to the commonly made assumption in reinforcement learning that the agent knows its own reward function. However, in practice, the brain would have to determine the correct weightings for these coefficients online, so as to best obtain homeostatic equilibrium. In practice, this requires some kind of control system to exist in the brain which computes the values of these coefficients depending on the physiological state and transmits these values to the dopaminergic system in the mid-brain. Such a control system is not modelled in this paper.

A simple example of the kind of control system that could be implemented, especially for maintaining homeostatic equilibrium, is PID control (Johnson & Moradi, 2005) around a homeostatic set-point. The central idea would be that the weights *θ*_*i*_ for a specific homeostatic variable would be set proportional to the deviation of that variable from an optimal set-point van Swieten and Bogacz (2020). Such a model is closely related to drive reduction theory (Hull, 1943; Juechems & Summerfield, 2019; Keramati & Gutkin, 2011) which fits closely with notions of homeostasis and allostasis in biology (allostasis here is achieved by updating long-term *values* based on the homeostasis of the weighting coefficients). Beyond simple homeostatic set-point controllers, it is possible that more complex cortical controllers also exist to achieve more fine-grained and context sensitive control over more abstract reward types such as social or monetary rewards.

## Methods

All experiments in this paper took place in tabular environments, where the state vector **x** is a one-hot vector over all possible states with a value of 1 for the state that the agent is in and 0 otherwise. The mathematical theory, however, is equally applicable to continuous-valued state vectors.

### Sea-Salt Experiment

In the simulated version of the experiments of (Robinson & Berridge, 2013) presented in Figure 3, the animals were exposed to the lever associated with the salt or the lever associated with the juice at random over 10 trials. If the salt lever was presented, the TD agent received a reward of −10 while if the juice lever was presented the agent received a value of +1. The TD agent learnt a value function with 2 states – juice and salt. The reward basis agent maintained separate salt and juice reward bases with the juice reward basis returning +1 for juice and 0 for salt and vice-versa for the salt basis. To match the rewards received, during testing in the deprivation conditions, we set *θ*_salt_ to +10 and maintained *θ*_juice_ to +1. The agents were trained over 10 trials with a learning rate of *η* = 0.1. Since there are no multi-step dependencies in this task, we trained with *γ* = 0.

The experimental data was replotted from Figure 3C in (Robinson & Berridge, 2013) including their error bars. To match the value functions of our agents to behavioural data, we assumed that the degree of appetitive approach and ‘liking’ behaviour plotted in the original experiment varied linearly to the value of the stimulus the animal had learned. That is, we simulated the simple equation,

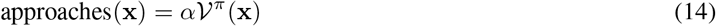

Where *α* is a free parameter of the model. We assume a different *α* scales approach for the food and juice levers. Hence each *α* parameter scales two values corresponding to approaching a given lever in two states. We fitted the *α* parameter to the homeostasis state and let the learner predict the salt-deprived state. The reward basis agent had additional *θ*_juice_ and *θ*_salt_ weighting coefficients which we fit instead of the *α* parameter. The TD agent cannot generalize because it possesses a single *α* parameter to fit both conditions which cannot vary due to physiological state, while the reward basis agent possesses two *θ* parameters, one for each state. For both agents we ran experiments over 10 seeds which resulted in slight variations in the value function due to the random presentations of either the salt or the juice lever.

### Predicting Striatal Dopamine

The reward basis agent was trained in a simulated version of the task paradigm used in (Papageorgiou et al., 2016) and graphically described in Figure 4A. The rats were randomly presented with a cue signifying either the food or sucrose lever was active. If they went to the cued lever, there was a 80% chance of getting that reward type, a 15% percent chance of switching to the other reward type (SWITCH condition) and a 5% percent chance of getting additional stimulus units (MORE condition). The reward basis agent maintained two reward and value bases over the two states of the experiment for food and sucrose respectively. The valuation and devaluation conditions were simulated by modifying the *θ*_food_ and the *θ*_sucrose_ physiological state weights used in the reward basis algorithm. The value functions were learned over 500 trials with 20 unique seeds. Differences in the experimental outcomes are due to random differences of the contingencies shown to the animal and hence slightly different value functions are learned. The dopaminergic responses are simulated as the sum of the weighted prediction errors upon stimulus presentation to the model, in accordance with the modulated prediction error model presented. That is, we identify,

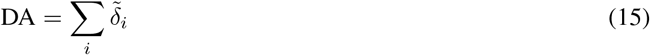

The experimental data is replotted from Papageorgiou et al. (2016). Figure 4B data comes from Figure 4a in Papageorgiou et al. (2016). We plot the relative dopamine level benchmarked against the dopamine level at lever extension where the ‘baseline’ level in (Papageorgiou et al., 2016) is normalized to 0.5 in our plot. That is, we set the level of dopamine reported as the ‘baseline’ in (Papageorgiou et al., 2016) to a value of 0.5 in our plot and transfer the dopamine levels reported in Papageorgiou et al. (2016) to our plot by measuring the deviations from this benchmark. The ‘valued’ experimental bar in our plot comes from the ‘satiation-valued’ dopamine level at the time of lever extension while the ‘devalued’ experimental bar comes from the ‘satiation devalued’ dopamine level at the time of lever extension. The free parameters in our model are the *θ*_*i*_ weighting coefficients which we fitted to the experimental data.

Experimental data in Figure 4C derives from Figures 2a and 2b in (Papageorgiou et al., 2016). The Dopamine experimental level is taken to be the dopamine level 7 seconds after reward delivery normalized against dopamine level at reward delivery – i.e. we plot the experimental dopamine level in our plot as the difference between the dopamine level at reward onset and 7 seconds after reward delivery in Figures 2a and 2b of Papageorgiou et al. (2016). This is why there are negative dopamine levels since this means that dopamine declined since reward delivery. Simulation results were generated by computing the prediction errors generated in case of an EXPECTED, MORE, or SWITCH condition. The *θ*_*i*_ weighting coefficients were free parameters which were fit to match experimental data by minimizing a least squares error between the model prediction given the parameter and the experimental value.

### Homeostatic Agent

Following Zhang et al. (2009), we implemented the ‘homeostatic agent’ which has also been proposed to explain zero-shot generalization in reward revaluation experiments. The homeostatic agent uses the standard TD update rule, to learn a cached representation of the value function but also, at test time, can dynamically modulate its estimate of learnt values dynamically using a ‘homeostatic’ term *κ* which modulates the predicted reward (but not the final function). Specifically, at test time, the homeostatic agent estimates a saliency-modified value function 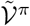 as,

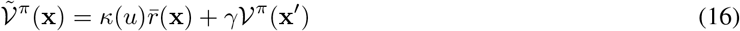

where *κ*(*u*) is a scalar parameter which is a function of the physiological state *u* and which scales the reward obtained in the value function update where 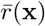 is the previously obtained reward in this contingency ^7^. The homeostatic agent can solve single-step tasks like the sea-salt experiment because it directly modulates the predicted reward from the outcome (salt or juice) according to the homeostatic state, however because it does not modulate the long-run value function, it cannot generalize to tasks with more than one step. We demonstrate that this is the case in the two-step task.

### Room Task

For the TD and reward basis agents, the value function was represented as a flattened vector of 6 × 6 = 36 states, which was initialized at 0. For the successor representation agent (see the next subsection for a detailed description of successor representations), the successor matrix was initialized as a 36 × 36 matrix of 0s and was updated on each timestep by the successor representation update rule (Equation 20). The successor agent computed its estimated value functions according to Equation 19.

For all agents a learning rate of *η* = 0.05, a discount factor of *γ* = 0.9 and a softmax temperature of 1 were used. Actions were selected by random sampling over the softmaxed distribution over actions. We used 500 steps between reversals. Means and standard deviations were obtained over 10 seeds for each agent in Figure 5.

### Successor Representations

The successor representation method in reinforcement learning allows for zero-shot reward revaluation by computing and caching the “successor” matrix which is the matrix of expected state transition probabilities. Given this matrix, it is possible to instantly recompute the value function if the reward function changes. The mathematics underlying the successor representation is extremely simple. Recall our matrix definition of the value function in terms of the transition operator *𝒯*.

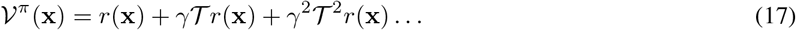

By a simple rebracketing, we can express this as,

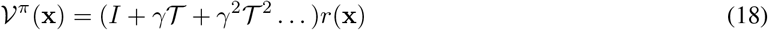

We can then separate out the matrix (*I* + *γ 𝒯* + *γ*^2^ *𝒯*^2^ …) and call it *ℳ*^*π*^, the successor matrix giving us the following equation for the value function (we use a *π* superscript to denote that it represents only the discounted state occupancies for a specific policy),

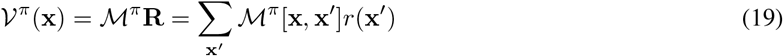

Where we use **R**(**x**) = [*r*(**x**_1_), *r*(**x**_2_) …] to denote the vector comprising the scalar reward for every possible state value, and the square brackets on *ℳ*^*π*^ [**x, x***′*] to denote indexing into the successor matrix *ℳ*^*π*^ at the state entries **x** and **x***′*. It is therefore clear that if we know *ℳ*^*π*^ and have it stored, then, given any change to the reward function **R**(**x**), it is trivial to recompute the correct value function *𝒱*^*π*^ (**x**) as a simple matrix multiplication of *ℳ*^*π*^ with **R**(**x**). This allows for instantaneous zero-shot reward revaluation since the value function can be recomputed so easily. The difficulty, of course, is in learning or optimizing the successor matrix *ℳ*. Fortunately, there is an iterative way of learning the successor matrix based on an estimate using only a single tuple of experience as with TD learning for the value function. The update rule is,

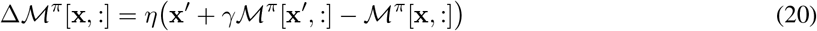

Where **x** is one-hot state vector which signifies the state of the agent before the transition, and **x***′* denotes the next state after the transition, *η* is a scalar learning rate parameter. The square brackets on *ℳ*^*π*^ denote indexing into a row of the successor matrix according to the state index – i.e., indexing *ℳ*^*π*^ [**x**, :] returns the row of estimated state occupancies of all possible state transitions **x***′* from the state before the transition **x**. Intuitively, the update rule updates the observed state from **x** to **x***′* transition by 1 while decaying the other non-observed state transitions to state **x***′* by *γ*. While this update rule is relatively straightforward and allows for easy and robust estimation of the successor matrix for discrete state spaces, a key computational imitation of the method is that the successor matrix *ℳ* is of dimension | *𝒳* | × | *𝒳* |, or the square of the size of the state-space. Thus, as the number of states grows, explicitly representing the full successor matrix becomes extremely memory intensive and eventually becomes intractable.

## Acknowledgements

The authors would like to thank Nathaniel Daw and Scott Waddell for discussion and Mycah Banks for her aid in preparing the figures. This work has been supported by BBSRC grant BB/S006338/1 and MRC grant MC_UU_00003/1.

## Code Availablility

Code to reproduce all the simulations and figures in this paper is freely available online at https://github.com/BerenMillidge/Reward_Bases.

## Appendix A: Additional Analyses of the Sea-Salt Experiment

Here we present results showing explicitly the choices and rewards obtained by agents in a variant of the Sea-Salt experiment, where the agent can select the lever on each trial. We chose to also simulate the bandit variant of the task so as to present a more detailed intuition about the behaviours and choices of agents under reward revaluation and more directly demonstrate the generalization performance of the reward basis algorithm, as well as the success of the homeostatic agent at generalizing in this task.

Instead of a Pavlovian paradigm where agents are simply presented with levers (which effectively serve as cues) during training and must learn the associations, here we instead simulate a two-armed bandit decision-making task where at each trial the rat must choose between a lever which dispenses sweet juice and always gives a reward of +1, and a lever which dispenses the sea-salt which gives a reward of −10 in the condition without salt-deprivation.

Upon reversal, we tested two conditions. In the *extinction* condition used originally in (Robinson & Berridge, 2013), after the reversal both levers did not give out their previous reward which we model in our simulation by setting the reward obtained from each lever to 0. Secondly, we also simulated the continuation of task in the salt deprived condition (*continuation condition*), where each lever continues to dispense its usual result after reversal, but now the salt lever in the salt-deprived condition gives a reward of +10. This condition allows us to compare the learning curves of the reward basis and TD learning agents and see that the reward basis agent achieves substantially faster generalization upon reward revaluation.

We tested three kinds of agents – TD learning agents, reward basis agents, and the homeostatic agent of (Zhang et al., 2009). All agents computed the value function using their respective methods. Then, actions were sampled randomly from a softmax distribution over computed Q values with a temperature parameter *β* equal to 1.

The reward basis agent uses two reward bases – one corresponding to receiving the juice (giving +1 when the juice was selected and 0 otherwise), and one corresponding to receiving the salt (giving +1 when the salt was selected and 0 otherwise). A set of reward bases like this seems plausible in the brain as the reward bases only represent basic tastes which can be distinguished by the hypothalamus and brainstem sensory regions such as the solitary nucleus, and which are of fundamental homeostatic importance. In the non-salt-deprived condition, the weightings for the reward bases were setup to mimic the standard reward structure so that *α*_*undeprived*_ = [1, −10] and in the salt-deprived condition the weighting coefficients were *α*_*deprived*_ = [1, 10]. For the homeostatic learner, the value of the *κ* homeostatic modulation term was set to 1 when in the non-salt-deprived condition and 10 in the salt-deprived condition.

All agents were initialized with a value function estimate of *𝒱*_0_ = [0, 0] and chose actions by sampling from an action posterior constructed by performing a softmax over the estimated value function of each state. This is because in this task there is a one-to-one correspondence between actions and states as each action simply moves the agent to that state. Below, we demonstrate the behaviours of our various agents on this simulated version of the sea-salt experiments. Actions were sampled randomly from a softmax distribution over computed Q values with a temperature parameter *β* equal to 1.

**Figure 6:**
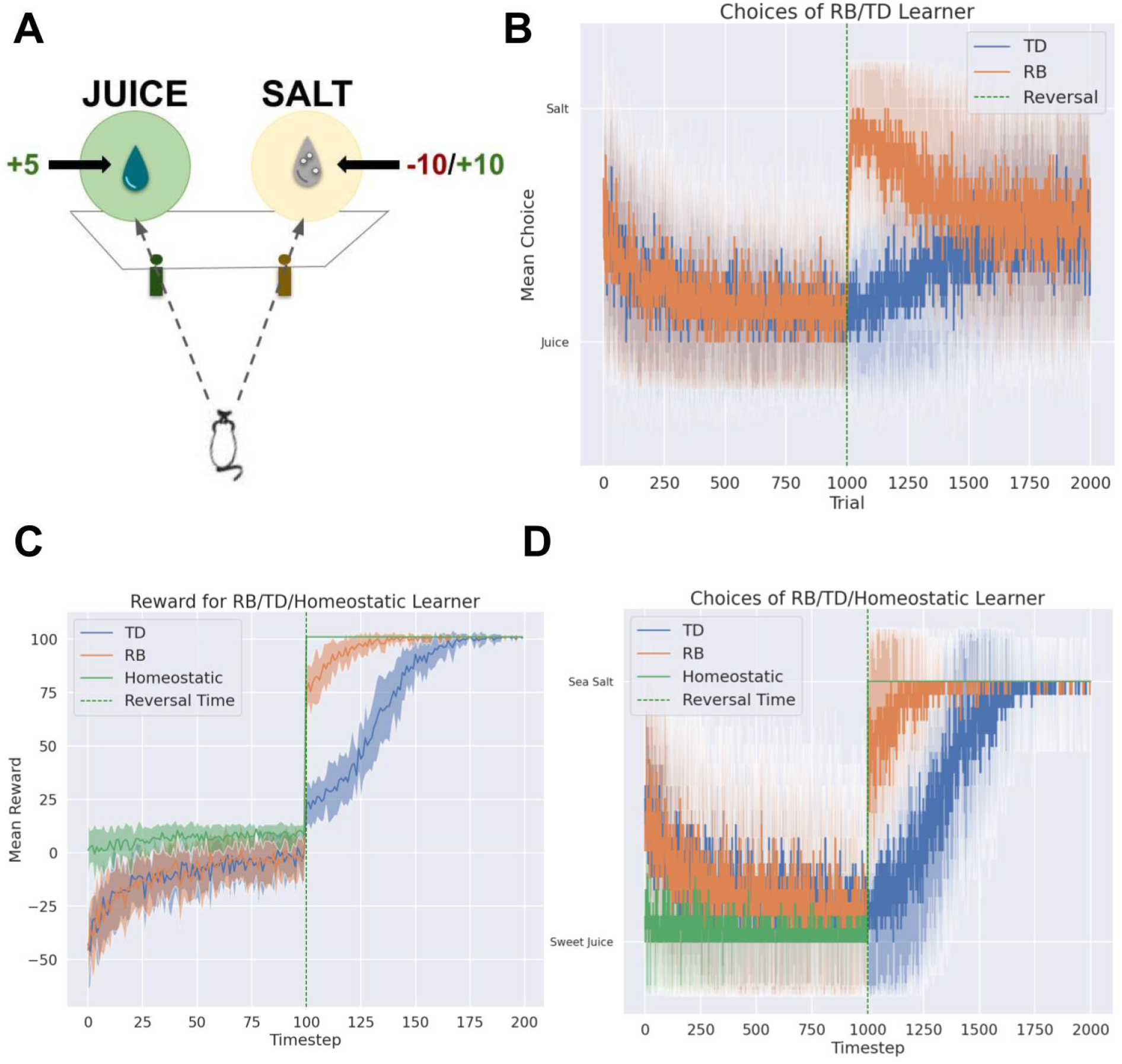
Simulations of bandit version of the salt deprivation experiment. **A**: The two-armed bandit variant of the Pavlovian task used in (Robinson & Berridge, 2013). The animal must choose between the two levers, one dispensing fruit juice and the other the salt solution. **B**: Choices made by the temporal difference (TD) and reward basis (RB) agent in extinction, where no reward for receiving the salt is ever given (both levers return 0 reward after reversal). The RB learner instantly switches to the salt even when it is presented in extinction, replicating the results of Robinson and Berridge (2013). **C** Rewards obtained by the reward basis (RB), TD and homeostatic learning agents in a bandit version of the task where the sea salt lever switches to dispensing positive reward, allowing TD learners to slowly learn the new contingencies. **D**. Choices made by the three agent types in a bandit version of the task. Presented are the mean values and standard deviations of rewards and choices computed over 20 seeds. The reversal occurred at timestep 100 out of 200. Agents were initialized with value function of zeros.

In the extinction condition, it is clear to see that the TD learner fails to generalize across the changing reward function induced by the salt deprivation state and continues to pull the sweet juice lever. On the other hand, the reward basis learner can instantly generalize and switch to the salt lever as soon as the reversal occurs, even though it has never experienced positive rewards from the salt lever, and continues with this behaviour for a substantial number of trials even though the salt lever is presented in extinction and never gives out any reward. Eventually, the choices of both agents in extinction decay towards a 50 − 50 split because after experiencing equal 0 rewards from both levers, the value function of each is slowly adjusted towards an identical 0 option, which results in random equiprobable choices between the two by the softmax decision rule used to select actions.

In the continuation condition, the TD learner fails to exhibit any generalization across changing reward functions and instead needs a length period of interacting with the task according to the new reward. This allows for the TD learner to eventually learn the correct contingency after repeated interactions. While the TD learner requires multiple interactions with the environment to learn, both the reward basis learner and homeostatic learner exhibit a clear zero-shot generalization capability, with both the reward obtained and the choices made abruptly reversing precisely at the reversal time without requiring any additional interactions with the environment. The generalization effect is especially pronounced for the homeostatic learner in that generalization is instantaneous and essentially perfect, while the reward basis learner shows considerable generalization immediately, but still requires some interactions with the environment to perform perfectly. This is due to the task essentially being one of reward estimation, which can be solved perfectly by simply estimating the rewards of each state correctly. The homeostatic learner has this ability essentially programmed in directly through the *κ* parameter, which is set so as to ensure that the predicted reward is precisely equal to the true reward of the environment. On the other hand, the reward basis learner was not essentially given direct access to the true reward, but only the change in the *θ* weighting coefficients – i.e. that the weighting for the salt had gone from −10 to 10. This then had to be combined with its value function estimate. However, this value estimate was derived from a TD learning algorithm on each reward basis, but since the salt was highly aversive initially, it was very rarely explored prior to the reversal leading to a poor estimate of its value compared to the juice option thus leading to a similarly poor estimate when the weighting coefficients changed. The reward basis approach thus allowed for a substantial degree of zero-shot generalization, but then additionally had to be finely tuned by more interactions with the environment. This highlights an inevitable limitation on the reward basis algorithm and comparable ones such as successor representations – namely that the value function in these algorithms is defined only for a fixed, non-optimal policy, and while the reward bases allow for the value function for a given policy to be changed instantaneously when the reward function is changed, changing the reward function almost always results in a different policy being optimal, and this change in the value function due to the change in the optimal policy is not captured by our method. The key advantage of our method compared to the homeostatic method is that it allows for the recomputation of the *value function* given a changing reward function, and is therefore able to dynamically recompute *multi-step behaviour* as the reward functions change rather than only recomputing greedy immediate reward maximization behaviour. We demonstrate this capability in the experiments on the two-step task.

## Appendix B: Additional Analyses of the Room Task

In this Appendix, we demonstrate that the reward basis agent can learn reward functions that are linear combinations of reward bases, as demonstrated by the mathematical results. Here, we varied the room task so that before the reversal only one of the three options was rewarded and the rest were punished. Then, after the reversal, the other *two* options were punished and the original one was punished. The reward function after reversal therefore consisted of a linear combination of the reward basis *r*(**x**)_after_reversal_ = 0 * *r*_1_(**x**) + 0.5 * *r*_2_(**x**) + 0.5 * *r*_3_(**x**). In Figure 7, we show that the reward basis agent can instantly generalize to this task as well, as can the successor agent due to the simpler nature of the task.

**Figure 7:**
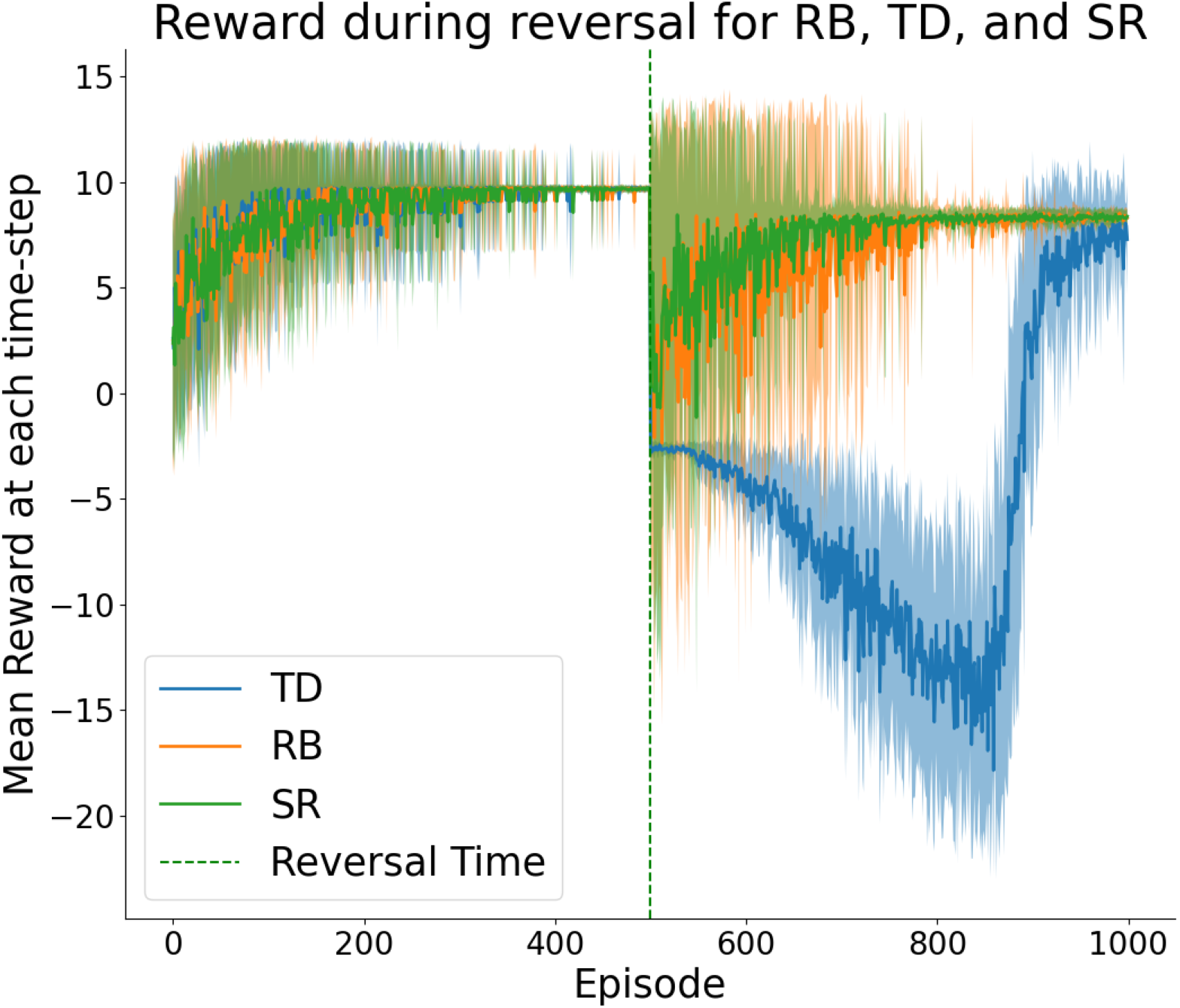
Instant generalization of the reward basis agent to a linear combination of reward functions, as predicted by the mathematical derivations. Both successor representation and reward basis agent show instant generalization while the TD agent must laboriously relearn the task.

## Footnotes

1 Mathematically, this difference can be represented by what the reward function is a function *of*. Standard reinforcement learning methods assume that the reward function is a function only of the state of the *environment*, but instead we can formalize it such that the reward function is a function of the state of the *agent*.

2 That TD-learning agents behave like this provides a clear evolutionary rationale for *not* using classic TD learning. If you are severely salt-deprived, it is not good enough to do what you always do until you randomly bump into the salt, and only then realize that it is more rewarding than expected. These situations instead require instant changes in behaviour *before* achieving any rewards.

3 Although this could perhaps be accomplished by a close interaction between the basal ganglia and hippocampus system which may encode successor representations (Stachenfeld, Botvinick, & Gershman, 2017)

4 With an infinitely long time horizon, everything works similarly but for simplicity, here we assume a finite time horizon *T*.

5 Since each reward and value basis can be computed independently in parallel, this method is well suited to parallel computation in the brain, where each reward basis could be computed using a separate physical neural circuit where only the results need to be combined at the end.

6 From a theoretical perspective this direct feedback connection is vital for DANs to be able to represent RPEs since a feedback connection is necessary for the DAN to compute the derivative of the value function (encoded in the MBON)

7 Another slightly different model also proposed by Zhang et al. (2009) is to define the reward term as log(*r*(*x*) + *κ*(*u*)). This approach only slightly changes the reward shaping and has no real impact on the fact that the homeostatic method, due to only modulating the reward, cannot perform zero-shot generalization across changing reward functions for multi-step tasks.

